# *In vitro* tau aggregation inducer molecules influence the effects of *MAPT* mutations on aggregation dynamics

**DOI:** 10.1101/2021.08.06.455436

**Authors:** David J. Ingham, Kelsey M. Hillyer, Madison J. McGuire, T. Chris Gamblin

## Abstract

Alzheimer’s disease (AD) and Alzheimer’s disease related dementias (ADRDs) affect 6 million Americans and they are projected to have an estimated health care cost of $355 billion for 2021. A histopathological hallmark of AD and many ADRDs is the aberrant intracellular accumulation of the microtubule associated protein tau. These neurodegenerative disorders that contain tau aggregates are collectively known as tauopathies and recent structural studies have shown that different tauopathies are characterized by different “strains” of tau filaments. In addition, mutations in the gene that encodes for tau protein expression have been associated with a group of tauopathies known as frontotemporal dementias with Parkinsonism linked to chromosome 17 (FTDP-17 or familial frontotemporal dementia). *In vitro* studies often use small molecules to induce tau aggregation as tau is extremely soluble and does not spontaneously aggregate in typical lab conditions and the use of authentic filaments to conduct *in vitro* studies is not feasible. This study highlights how different inducer molecules can have fundamental disparities to how disease related mutations effect the aggregation dynamics of tau. Using three different classes of tau aggregation inducer molecules we characterized disease relevant mutations in tau’s PGGG motifs at positions P301S, P332S, and P364S. When comparing these mutations to wild type tau, we found that depending on the type of inducer molecule used we saw fundamental differences in total aggregation, aggregation kinetics, immunoreactivity, and filament morphology. These data support the hypothesis that different tau aggregation inducer molecules make different polymorphs and perhaps structurally distinct strains.

## Introduction

Neurodegenerative disorders are often characterized by the aggregation of one or more proteins^1^. In Alzheimer’s disease (AD) and Alzheimer’s disease related dementias (ADRDs), the microtubule associated protein tau accumulates within neurons and glia of the central nervous system. These terminal maladies are not only devastating to the 6.2 million Americans who suffer from them, but also cause patients to require round the clock care during advanced stages of disease. This effect is felt more broadly by society as AD and ADRDs are estimated to have associated health care costs of $355 billion in the United States for 2021 and an estimated 11 million unpaid caregivers^2^. To make matters worse, the number of cases and associated costs of AD and ADRDs are expected to rise dramatically over the next few decades.

The aberrant accumulation of tau into beta sheet enriched amyloid folds correlates strongly with the progression and severity of cognitive decline in AD patients^3^. In AD, tau primarily accumulates into twisted paired helical filaments (PHFs) and untwisted straight filaments (SFs). Other tauopathies can include PHFs or SFs, but many are characterized by tau filaments dissimilar to those found in AD. ADRDs include Pick’s disease, progressive supranuclear palsy, corticobasal degeneration, chronic traumatic encephalopy, as well as other frontotemporal dementias with Parkinsonism linked to chromosome 17 (FTDP-17, or familial frontotemporal dementias - fFTD). FTDP-17 tauopathies are of particular interest to the research field because as well as having tau accumulation as a histopathological hallmark, they have been associated with over 50 different intronic and exonic mutations of the *MAPT* gene that encodes the expression of all six isoforms of tau in the human adult central nervous system^4^.

The nomenclature of the 6 tau isoforms expressed in adults is based on the inclusion of 0, 1, or 2 N terminal domains, as well as the inclusion of 3 or 4 microtubule binding repeat domains (MTBR). This results is the 6 tau isoforms of the central nervous system being named 2N4R, 1N4R, 0N4R, 2N3R, 1N3R, or 0N3R^5^. Each of the microtubule binding repeats ends with a PGGG motif. Interestingly, a P to S substitution mutation on three of the four PGGG motifs has been associated with cases of FTDP-17 at positions 301^6^, 332^7^, 364^8^ (numbering based on full length 2N4R human tau isoform). In addition, P301S is one of the most common mutations used in both *in vitro* and *in vivo* tau aggregation model systems, primarily due to the formation of PHF-like filaments, pro-aggregation properties, and relatively poor affinity towards microtubules^9^. The PGGG motif found at the end of microtubule binding repeat 1, position 270, has not been associated with disease linked mutations. Although recent structural studies of tau filaments isolated from disease have shown that this region of tau, MTBR 1, does form part of the ordered filament core isolated from the three repeat tauopathy [Pick’s disease (PiD)^10^], it is not found as part of the ordered fibril core of mixed 3R-4R tauopathies [AD^11^ and chronic traumatic encephalopathy (CTE)^12^], as well as the 4R tauopathy, [corticobasal degeneration (CBD)^13^].

In this study, we compare the aggregation characteristics of three of these FTDP-17 P to S mutations, as well as the non-disease related P270S mutation, to WT 2N4R tau. We used site directed mutagenesis to recombinantly express and purify each of the P to S mutations at positions 270, 301, 332, and 364 in the full-length isoform of human tau, 2N4R (HT40) (supplemental figure S1).

However, because tau is natively unfolded, contains high numbers of both positively and negatively charged residues, and is highly soluble in solution, it is resistant to spontaneous aggregation^5^. Therefore, biochemical “inducers” of tau aggregation are widely employed to initiate and enhance the aggregation of tau *in vitro*. One of the most commonly used tau aggregation inducers, heparin^14^, induces polymorphic tau aggregate structures that are dissimilar to any structures found in filaments isolated from disease^14, 15^. Heparin is therefore not likely to be a useful model in studies characterizing and identifying tau aggregation-based therapeutics or the molecular dynamics of aggregation. Therefore, we chose three alternative inducers of tau aggregation for this study: the polyunsaturated fatty acid arachidonic acid (ARA), polyphosphate, and RNA, although to date there have not been any high-resolution structures published of *in vitro* tau filaments generated with these inducers.

We have previously found that ARA rapidly polymerizes tau to form filaments that have similar morphological characteristics to straight filaments isolated from AD^16^. In addition, ARA is found within the intracellular environment at elevated levels during times of oxidative stress and could play numerous roles in the pathology of AD^17^. Furthermore, antibodies raised against ARA induced filaments have been shown to have a high affinity towards aggregated tau in diseased brain tissue^18, 19^. We have also shown that two different small molecule tau aggregation inhibitors (TAIs), the isoquinoline ANTC-15 and the phenothiazine LMTX, appear to inhibit heparin and ARA induced filaments in an inducer-specific manner^20^. For example, ANTC-15 inhibits ARA induced filaments, but not heparin induced filaments. Conversely, LMTX inhibits heparin induced filaments, but not those induced by ARA. It is therefore likely that the polymorphs formed from ARA and heparin induction are structurally distinct. Polyphosphate is present in mammalian neurons and has been shown to induce the aggregation of tau *in vitro*^21–23^. RNA has been shown to induce the aggregation of tau *in vitro* ^24, 25^ and tau aggregates in disease can sequester RNA^26^. Although the structures formed by ARA, polyP, and RNA are not known and it is unclear whether they play a direct role in tau aggregation in disease progression, they have the potential to form biologically relevant, and potentially disease relevant, aggregates of tau.

Using right-angle laser light scattering, transmission electron microscopy, and conformationally sensitive ELISA assays, we compared the maximum protein aggregation, filament length, morphology, and immuno-reactivity of toxic tau species formed *in vitro* by wild type tau and tau variants in the presence of different classes of inducers and different sizes of inducers within a class.

To our knowledge, not only is this the first study to complete a direct biochemical comparison of this group of disease related mutations, it is also the first to directly compare multiple *in vitro* aggregation inducers to study biochemical characteristics of multiple disease related mutations. Using this combination of approaches we have found that different classes of tau aggregation inducer molecules can not only influence typical aggregation characteristics such as length of filaments and total amount and rate of aggregation, but also, the type of inducer used can have effects on the fundamental differences between wild type tau and mutant constructs as well as immunoreactivity towards conformationally sensitive antibodies. The data strongly supports the hypothesis that filaments formed in the presence of different inducer molecules have different characteristics in terms of aggregation morphology, aggregation dynamics, assay compatibility, and immuno-reactivity. These findings illustrate the importance of identifying disease relevant inducer molecules to be used in studies of characterizing disease related mutations.

## Materials and Methods

Chemicals and reagents: Full length 2N4R tau (HT40, 441 amino acids) and all mutant constructs were expressed and purified as previously described^27^. Using the HT40 Pt7c WT construct, amino acid substitutions were introduced using a QuikChange II XL site directed mutagenesis kit (200521) purchased from Agilent (Santa Clara, CA). After transformation into *BL21-Gold (DE3)* competent cells, protein was expressed and purified using Ni-His Tag affinity purification and size exclusion chromatography. King et. al. has shown that the poly-histidine tag does not appear to influence tau aggregation and therefore was not removed prior to concentration quantification and subsequent *in vitro* studies. The concentration of protein was quantified using a Pierce BCA protein assay kit (23225) purchased from Thermo Fisher Scientific (Rockford, IL), each protein prep was at a concentration between 1 and 2 mg/mL. Tau purity and concentration was confirmed by SDS PAGE. Individual aliquots of 50-100 µL were prepared and stored at −80°C and a fresh aliquot was used for each experiment to avoid repeated freeze/thaw cycles. Arachidonic acid (90010) was purchased from Cayman Chemical, (Ann Arbor, MI). Polyphosphate medium chain – [P100 (EUI005) - a heterogenous mixture with most chains being between 45-160 phosphate units and a purity of <1% monophosphate] and long chain - [P700 (EUI002) – a heterogenous mixture with most chains between 200 and 1300 phosphate units and a purity of <1% monophosphate] was purchased from Kerafast (Boston, MA). TOC1, TNT1, and Tau 5, Tau 7, and Tau 12 antibodies were a kind gift from Dr. Nicholas Kanaan, Michigan State University. Each of these antibodies was at a concentration of approximately 1 mg/mL. T22 antibody (ABN454) was purchased from Millipore Sigma, (Burlington, MA). The primary detection antibody (Tau 5, 7, and 12 were used as primary detection against T22 capture antibody) was an anti-tau polyclonal rabbit antibody (A002401-2) purchased from Agilent (Santa Clara, CA). A goat anti-rabbit IgG (H+L) and goat anti-mouse IgG (H+L) antibody with HRP conjugate (1706515 and 1706516, respectively, Bio-Rad, Hercules, CA) was used as a secondary detection antibody. Qiagen miRNeasy mini kit (217004) and Qiagen RNeasy minielute clean up kit (74204) was purchased from Qiagen (Germantown, MD). HEK293T cells (ATCC CRL-3216) were kindly provided by Dr. David Davido, University of Kansas. Mini Trans-Blot pre-cut filter paper (1703932) and 0.22 µm nitrocellulose membrane (16020112) was purchased from Bio-Rad (Hercules, CA). Chemiluminescent kit was a Supersignal West pico plus chemiluminescent substrate (34577) purchased from Thermo Scientific (Rockford, IL).

RNA Isolation: Mammalian RNA was isolated from HEK293t cells using 2 different procedures. HEK 293T (ATCC CRL-3216) cells were maintained in Dulbecco’s Modified Eagle’s Medium (Cytiva) and supplemented with 5% fetal bovine serum (FBS), 2mM L-glutamine, 10 U/mL penicillin, 10 U/mL Streptomycin. Cells were grown in BioLite 175 cm^2^ vented flask (Thermo Scientific) and maintained in a humidified incubator containing 5% CO_2_ at 37 °C.

To isolate the small RNA (RNA <200 nts) and long RNA (RNA >200 nts) separately, a modified version of the Qiagen miRNeasy mini kit and minielute cleanup kit isolation procedure was used to isolate samples into small RNA or long RNA fractions. The cells were lysed using QIAzol Lysis Reagent by adding 8.75 mL to the cell-culture dish. The lysate was collected and vortexed to mix, then stored in 700 μL aliquots at −80 °C. After thawing, the homogenate was incubated at room temperature (∼20 °C) for 5 minutes. Under a fume hood, 140 μL of chloroform was added to the tube containing the homogenate and vortexed vigorously for 15 s. The tube was incubated at room temperature for 2-3 minutes and then centrifuged for 15 minutes at 12,000 × g at 4 °C. The upper aqueous phase was transferred to a new collection tube and 1 volume of 70% ethanol was added and mixed thoroughly by vortexing. The sample was pipetted into a RNeasy Mini spin column placed in a 2 mL collection tube and centrifuged at 10,000 × g for 15 s at room temperature (15–25 °C). The flow-through was pipetted into a 2 mL reaction tube. The used spin column was set aside to isolate long RNA. 450 μL of 100% ethanol was added to the flow-through and mixed thoroughly by vortexing. 700 μL of the sample was pipetted into a RNeasy MinElute spin column placed in a 2 ml collection tube, then centrifuged for 15 s at 10,000 × g at room temperature. The flow-through was then discarded and this was repeated until the whole sample had been pipetted into the spin column. 500 μL Buffer RPE was then pipetted into the RNeasy MinElute spin column and centrifuged for 15 s at 10,000 × g, the flow-through being discarded. Next, 500 μL of 80% ethanol was added to the RNeasy MinElute spin column and centrifuged for 2 min at 10,000 × g to dry the spin column membrane. The flow-through and collection tube were discarded. The spin column was placed into a new 2 mL collection tube and centrifuged for 5 min at 10,000 × g. The RNeasy MinElute spin column was then placed into a 1.5 mL collection tube and 14 μL RNase-free water was pipetted onto the spin column membrane. It was then centrifuge for 1 min at 10,000 × g to elute the small RNA-enriched fraction.

Using the previously reserved RNeasy Mini spin column, the long RNA was eluted. 700 μL of Buffer RWT was added into the RNeasy Mini spin column and centrifuged for 15 s at 10,000 × g to wash the spin column membrane. The flow-through was discarded and 500 μL of Buffer RPE was added to the RNeasy Mini spin column. It was centrifuged for 15 s at 10,000 × g and the flow-through was discarded. Another 500 μL of Buffer RPE was added into the RNeasy Mini spin column and centrifuged for 15 s at 10,000 × g. The flow-through and collection tube were discarded. The RNeasy Mini spin column was placed in a new 2 mL collection tube and centrifuged at16,000 × g for 1 min. The RNeasy Mini spin column was then placed into a new 1.5 mL collection tube and 30 μL of RNase-free water was pipetted directly onto the spin column membrane. It was centrifuged for 1 min at 10,000 × g to elute the total RNA. This process was repeated for all samples of HEK293t cells. After this protocol was completed, a high sensitivity RNA TapeStation was used to run 2 μL samples of both fractions of RNA to confirm the size fractioning, the results of which showed significant size fractioning was achieved (figure S2). In order to determine the concentration of the differing RNA samples, readings were taken using a nanodrop and diluted to a concentration of 270 ng/μL using RNase-free water. The samples were stored at −80 °C until used in reactions.

Aggregation Reactions: Reactions were set up using each of the P to S tau mutations, P270S, P301S, P332S, P364S, as well as wild type (WT) tau. A no tau inducer only negative control and no inducer tau only negative control was also prepared and incubated with all other samples. Each mutant and WT tau were induced with small RNA, long RNA, P100, P700, or ARA in separate reaction tubes. All reactions were performed in 1.5 mL reaction tubes with a total volume of 200 μL. Ultrapurified molecular biology grade H_2_O was added first, followed by 4 μl of 250 mM DTT to the reaction tube and mixed by pipetting and lightly tapping the reaction tube. 1 M NaCl was added to bring the final NaCl concentration to 100 mM for ARA reactions or 25 mM for polyphosphate and RNA reactions. Again, each sample was mixed by pipetting and gentle tapping. HEPES at a pH of 7.64 was added in 8 μL volume of 250 mM to a final concentration of 10 mM. After mixing by pipetting and gentle tapping 20 μL of 1 mM EDTA stock was added in the same manner for a final concentration of 0.1 mM.

To ensure RNase activity did not degrade the RNA inducer, a stock of EDTA, HEPES, NaCl, and DTT was also made using DEPC treated H_2_O. However, there was no significant changes to RNA induced aggregation of WT tau using DEPC treated reagents when compared to preliminary studies that did not use DPEC treated H_2_O (data not shown). Either WT or mutated tau was then added to a final concentration 2 μM and mixed by pipetting and gently tapping.

Inducer was then added as follows to the respective samples, 10 μL of either small RNA or long RNA was added for a final RNA concentration of 13.5 ng/μL, 10 μL of either P100 or P700 was added for a final concentration of 10 ng/μL, and 7.5 μL of 2 mM ARA diluted in 100% ethanol was added to give a final concentration of 75 μM ARA 3.75% ethanol. For the controls, a Sup200 buffer (250 mM NaCl, 10 mM HEPES, 0.1 mM EGTA, pH 7.64) was used in place of the tau and RNase-free water was used in place of the RNA inducer, molecular biology grade H_2_O was used for no polyphosphate control and 7.5 μL of ethanol for the no ARA control. The reaction tubes were then incubated without agitation at 37 °C for 72 hours for RNA, 48 hours at 37 °C for polyphosphate, and 20 hours at 25 °C for ARA.

Sandwich ELISA: Following aggregation reactions, samples were analyzed using a modified sandwich ELISA assay based on previously described conditions^20, 28^. Capture antibody was used to coat Corning 3590 EIA/RIA 96 well microplate wells at a volume of 100 µL/well of either TOC1 (2 ng/µL), TNT1 (1 ng/µL), T22 (1ng/µL), or a mixture of Tau 5, Tau 7, and Tau 12 antibodies (referred to as 5,7,12) (1 ng/µL each). Capture antibodies were diluted in BSB capture buffer (100 mM boric acid, 25 mM sodium tetraborate, 75 mM NaCl, 250 µM thimerosal, pH 8.56). Plates were then sealed and incubated with gentle agitation overnight at 4 °C. After capture antibody incubation, the plate was blotted and washed 2x with 300 µL/well of BSB wash buffer (100 mM boric acid, 25 mM sodium tetraborate, 75 mM NaCl, 250 µM thimerosal, 60 mM BSA, 0.1% Tween 20, pH 8.56). Plates were then blocked and incubated for a further 1.5 hours with 300 µL of 5% non-fat dry milk (NFDM) dissolved in BSB wash buffer sealed at room temperature with gentle agitation. Samples were diluted in 5% NFDM BSB wash buffer to a concentration of 100 nM for TOC1 capture antibody, 25 nM for TNT1, 50 nM for T22, and 50 nM for 5,7,12. To provide an internal standard curve, dilution series of no compound polymer and monomer controls were added to the plate at a range of (3.125 nM - 400 nM for TOC1, 3.125 nM - 75 nM for TNT, and 1.5 nM-150 nM for T22). In our hands, the EC_50_ of the polymerized tau affinity curve was found to be 105 nM, 28 nM, and 35 nM for TOC1, TNT1, and T22 respectively. As 5,7,12 detects total tau, only a monomer standard curve was used at dilutions of 5-200 nM. Samples were added to a volume of 100 µL/well. Plates were sealed and incubated with gentle agitation for 1.5 hours at room temperature. Following incubation, plates were washed 2× using BSB wash buffer. A primary detection antibody was added at volumes of 100 µL/well. For TNT1, TOC1, and 5,7,12, polyclonal rabbit detection antibody diluted to a concentration of 50 ng/mL in 5% NFDM BSB wash buffer was added. For

T22 capture antibody, 5,7,12 was added at a concentration of 1:1,000 dilution. Further incubation was carried out after sealing the plate at room temperature for 1.5 hours with gentle agitation. Following incubation with primary detection antibody, plates were washed 2× using BSB wash buffer before the addition of appropriate secondary detection antibody (100 µL/well of the goat anti-rabbit IgG for TOC1, TNT1, and 5,7,12 capture antibody, and 100 µL/well of goat anti-mouse IgG for T22 capture antibody). Both secondary detection antibodies were diluted 1:5,000 in 5% NFDM BSB wash buffer. The plate was sealed and incubated at room temperature with gentle agitation for 1.5 hours. After incubation, plates were washed 3× using BSB wash buffer before the addition of 50 µL per well of tetramethylbenzidine (TMB) substrate. Plates were then covered and incubated with gentle agitation at room temperature for 20 minutes before the addition of 50 µL of a 3.6% H_2_SO_4_ stop solution. Readings were taken at an absorbance of 450 nm using a Varian Cary 50 UV Vis spectrophotometer with a Varian Cary microplate reader. Raw data readings were zeroed against a monomeric control of each mutant and then converted to % light absorbance. In the case of 5,7,12 capture antibody ELISA, the polymerized reactions were normalized against the monomeric protein for each mutation and WT. Statistical analyses were completed using an un-paired t-test to compare each mutation to WT 2N4R for TOC1, TNT1, and T22 ELISAs. In the case of 5,7,12, a Tukey’s multiple test was completed. For both tests statistical significance was defined as ∗ *p* ≤ 0.05; ∗∗ *p* ≤ 0.01; ∗∗∗ *p* ≤ 0.001.

For this experiment four different capture antibodies were used, 5,7,12 (a mixture of 3 monoclonal total tau antibodies that bind to residues 9-18 tau-12^29^, 218-225 tau-5, and 430-441 tau-7^30^,), TNT1^18^ (binds to the phosphatase activating domain epitope at residues 7-12 that are made accessible through tau fibrilization), TOC1^19^ (recognizes an epitope between residues 209- 224 with a high affinity for small tau oligomers and larger aggregates), T22^31^ (has been shown to bind specifically to tau oligomers that have been seeded using Aβ42 oligomers as well as *in vitro* heparin induced oligomers).

Transmission electron microscopy: Samples were diluted 1:10 in polymerization buffer and fixed with 2% glutaraldehyde for 5 minutes at room temperature. The samples were then affixed to a 300-mesh carbon formvar coated copper grid, purchased from Electron Microscopy Sciences, (Hatfield, PA) by floating the grid on a 10 µL droplet of sample for 1 minute. The grid was then blotted on filter paper and washed on a droplet of ddH_2_O before being blotted and stained by floating the grid on a droplet of 2% uranyl acetate as previously described^32^. Each grid was imaged using a JEOL JEM 1400 transmission electron microscope fitted with a LaB_6_ electron source (Electron Microscopy Research Lab, University of Kansas Medical Center). Five random images per grid were taken at a 5,000× magnification (to improve statistical power, 15 images were taken for both small RNA and long RNA induced filaments. Images were analyzed using Image Pro Plus 6.0 software by measuring the number, length, area, and perimeter of filaments >25 nm in length. Under our experimental conditions, it is very difficult to identify filaments less than 25 nm. To avoid erroneous results the assay has been limited to measuring tau filaments and oligomers greater than 25 nm.

Right-angle laser light scattering: Aggregation reactions were analyzed using right-angle laser light scattering as previously described^33^ to determine the amount of aggregated material. The average light intensity measured of each sample was zeroed against a no inducer monomeric control for the respective tau mutant being imaged, and a no tau/inducer-only control by subtracting the background signal from the measured signal of the endpoint aggregation reactions. Briefly, samples were transferred to a 5 mm × 5 mm optical glass fluorometer cuvette (Starna Cells, Atascadero, CA) in the light path of a 532 nm wavelength 12 mW solid-state laser operating at 7.6 mW (B&W Tek Inc. Newark, DE) and images were captured using a Sony XC- ST270 digital camera with an aperture of f/ 5.6. Images were analyzed using Adobe Photo Shop 2021 by taking histogram readings of the pixel intensity across the scattered light path.

Right-angle laser light scattering kinetics of aggregation: Using the right-angle laser light scattering assay described above, samples were placed into a cuvette at time zero prior to the addition of the respective inducer molecule. An image was captured prior to induction and at time 0 immediately after induction of each protein with either ARA, P100 or P700 as inducers. In the case of ARA, readings were taken every 5 minutes between 0- and 30- minutes post induction (p.i.), then at 45, 60, 90 and 120 minutes p.i. then every hour until 6 hours p.i. In the case of P700, images were taken at subsequent time points every 5 minutes p.i. for the first 60 minutes p.i. then at the following p.i. time points: 1.5hrs, 2hrs, 3hrs, 4hrs, 6hrs, 8.5hrs, 10hrs 40 minutes, 15hrs, and 16hrs. In the case of P100, images were taken at the same timepoint p.i. as for P700 (except 16hrs p.i.) with additional readings at 24 and 48 hours.

Thioflavin fluorescence: A standard assay in tau aggregation studies is thioflavin fluorescence using thioflavin S or thioflavin T. While thioflavin fluorescence is a useful tool in monitoring ARA and polyphosphate induced filament formation, we found that RNA gave a false positive result when using thioflavin T and quenched fluorescence of thioflavin S (data not shown).

Dot-blot assay: A 0.22 µm nitrocellulose membrane was pre-soaked for 10 minutes in Tris Buffered Saline (TBS – 500 mM NaCl, 20 mM Tris, pH 7.5). Polymerized 2N4R tau samples were diluted to a concentration of 20 ng/uL in TBS and added to the membrane using a dot-blot manifold (no vacuum). Samples were incubated on the membrane for 30 minutes at room temperature before removal of excess liquid and blocking the membrane with 5% non-fat dried milk (NFDM) in TBST (TBS + 0.05% Tween 20). The membrane was blocked for 1.5 hours with gentle agitation at room temperature. After incubation, the membrane was washed for 5 minutes 3× with TBST. The T22 primary detection antibody was diluted at 1:1,000 concentration in 5% NFDM in TBST, the membrane was submerged in T22 and incubated at room temperature for 2 hours with gentle agitation. Secondary antibody (Goat anti-rabbit IgG) was diluted in 5% NFDM in TBST to a 1:3,000 concentration. The membrane was washed 2× in TBST, before being submerged in secondary antibody at room temperature for 2 hours with gentle agitation. The membrane was once again washed for 5 minutes 2× using TBST before being developed using Thermo Fisher Supersignal West Pico Plus chemiluminescent substrate (catalog number 34580). An image of the blot was taken using UVP Chemidoc IT_2_ western blot imager and analyzed using Adobe Photoshop software using the histogram function to measure dot-blot intensity (figure S5).

## Results

### ARA induced tau aggregation

Using ARA as an inducer molecule, we initially completed LLS and TEM endpoint studies by allowing aggregation experiments to run for 20 hours (figure 1). The results from the LLS experiments (figure 1A) revealed that the P301S mutation caused significantly more aggregation when compared to WT. In addition, P332S led to a significant decrease in aggregation when compared to WT. Although not statistically significant, the average total amount of aggregation of the P270S mutation appeared to decrease (approximately 15% less than WT), and P364S showed no change when compared to WT.

**Figure 1:**
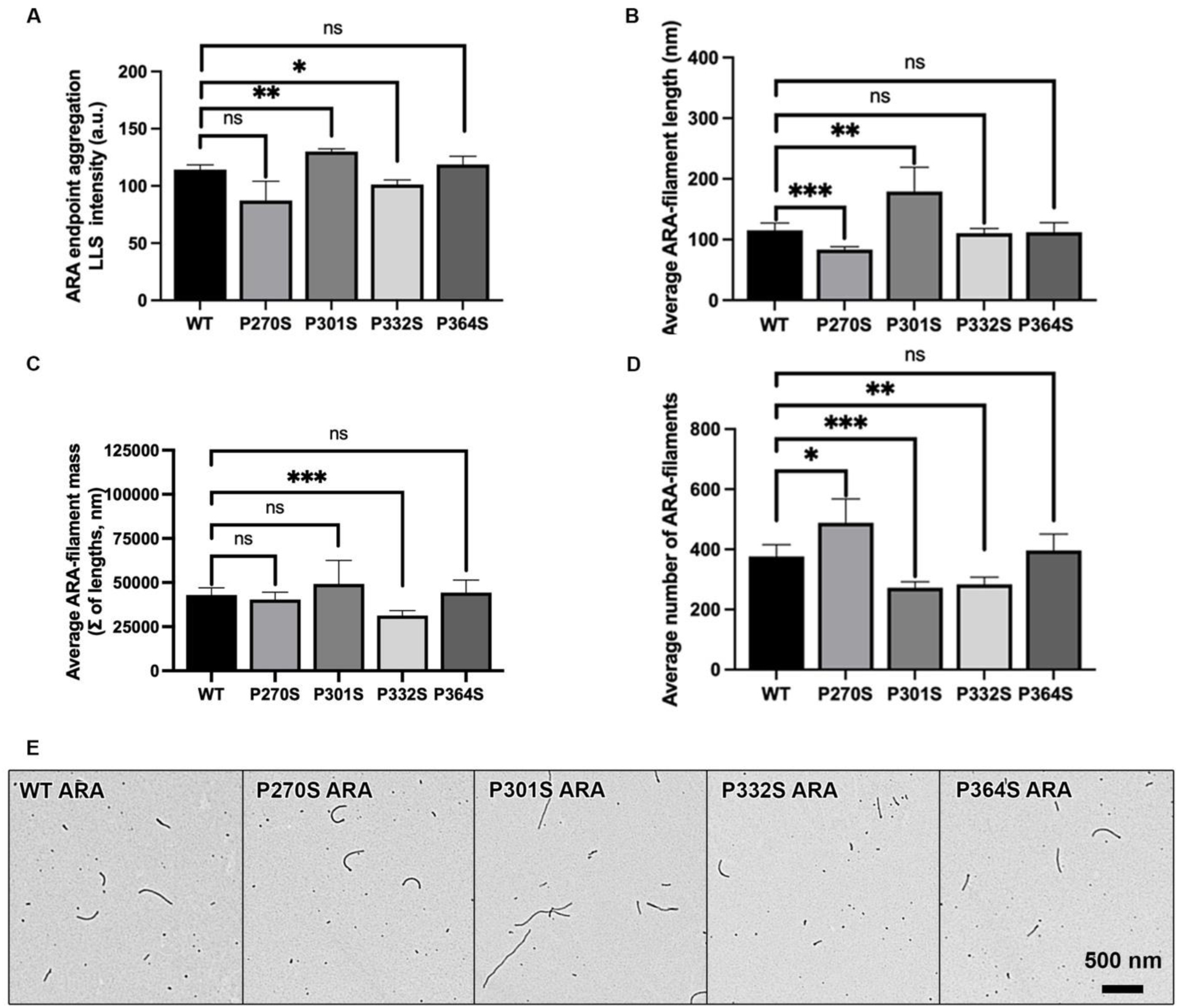
LLS and TEM endpoint measurements of ARA aggregation reactions. A) Endpoint total amount of aggregation of ARA induced aggregation for WT and each of the four P to S mutations, quantified using laser light scattering (n=3 ± s.d.). 5 different TEM images selected at random were quantified to measure. B) average ARA induced filament length (>25 nm) ± s.d., C) average ARA filament total filament mass of each micrograph ± s.d., and D) average number of filaments (>25 nm) per micrograph ± s.d.. E) Representative TEM micrographs of WT and each of the four P to S mutations (see labeling in upper left corner of each micrograph). Scale bar in bottom right corner of P364S image represents 500 nm and applies to each micrograph. Error bars on figures A-D represents SD of each data set. Each P to S mutation in A-D was compared to WT using an unpaired t-test. *ns p* > 0.05, ∗ *p* ≤ 0.05; ∗∗ *p* ≤ 0.01; ∗∗∗ *p* ≤ 0.001. *ns* = not significant.

Quantitative TEM was used to measure average filaments length (figure 1B), total filament mass (figure 1C), and number of filaments (figure 1D). Consistent with the findings of the LLS experiments, the difference in filament mass of P270S filaments was not statistically significant when compared to WT. However, the P270S mutation caused a greater average number of filaments that were shorter than WT filaments. Consistent with results from other studies, P301S filaments induced by ARA were significantly longer than WT filaments, however, there were also fewer P301S fibrils. This resulted in P301S having a higher average mass per image, but not enough to reach the threshold of statistical significance (p-value < 0.05). No significant difference in average filament length was seen when comparing WT to the P332S mutation, however there were fewer filaments that resulted in a significant decrease in filament mass, consistent with the LLS results. The P364S mutation cause no significant difference in filament length, mass, or number of filaments when compared to WT tau.

### ARA induced aggregation measured by sandwich ELISA

The 5,7,12 ELISA showed no significant difference between WT and any of the P to S mutations (figure 2A). This shows that the mutations themselves do not appear to significantly inhibit the affinity of the two total tau antibodies; either polyclonal rabbit or 5,7,12 (figure 2A). In contrast with the LLS results, there was no significant difference between ARA induced WT tau and any of the mutations when detecting with either TNT1 or TOC1 antibody (figure 2 B-C). However, the TOC1 ELISA did show that the average P301S signal was higher than WT, but not enough to meet the pre-determined p-value threshold of 0.05. Based on the LLS results we would have expected the TNT1 ELISA to show an increase for P301S and a decrease for P332S. Interestingly, ARA induced P301S, P332S, and P364S had significantly less reactivity with the T22 antibody than ARA induced WT tau, however P270S was not significantly different (figure 2D).

**Figure 2:**
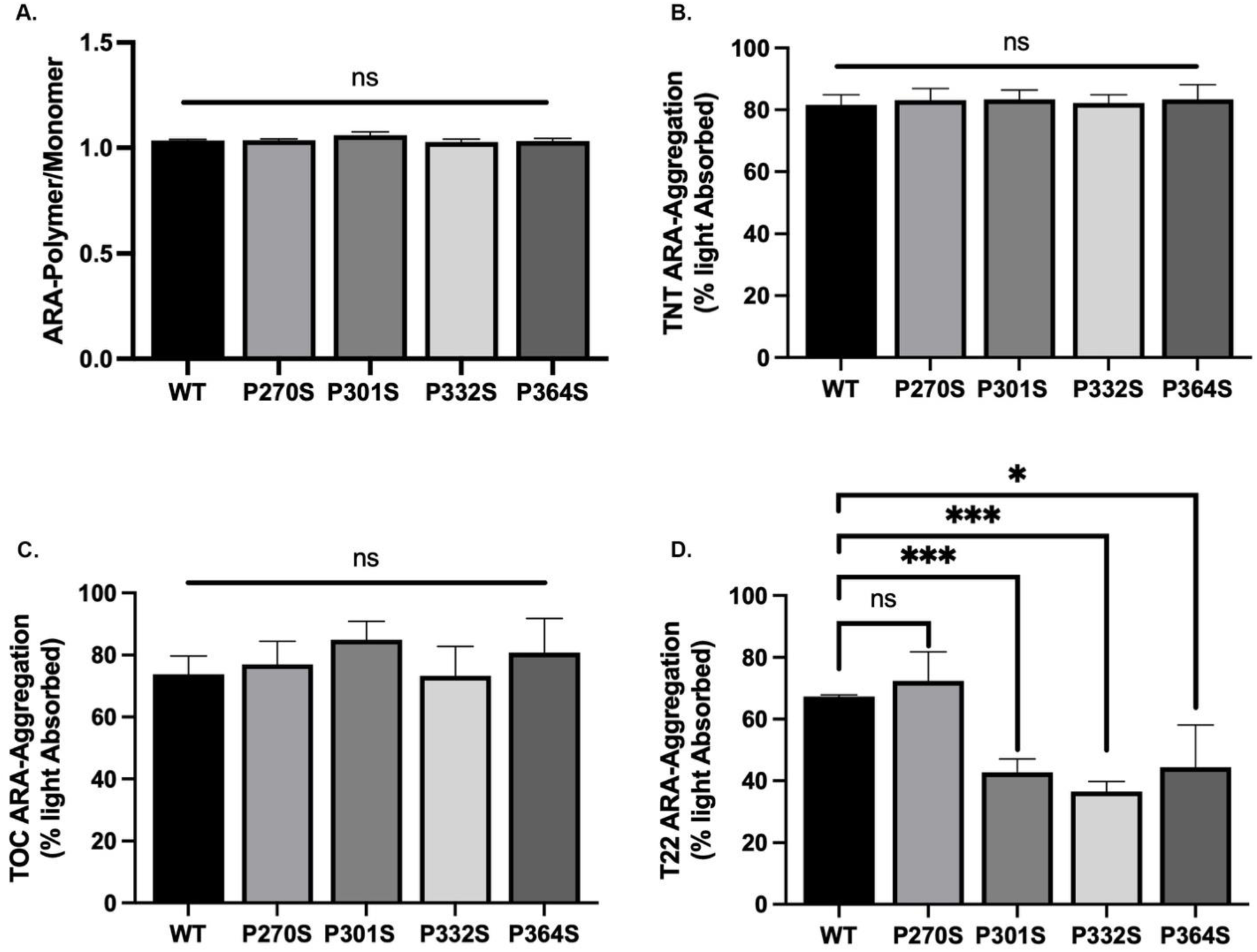
ARA ELISA data. ELISA results using 5,7,12 monoclonal total-tau antibody mixture as a capture antibody. The signal for each aggregate was divided by the signal for monomeric tau (either WT or the respective P to S mutations). Y axis represents a fraction of monomeric tau signal (e.g. 1=100%). B-D, ELISA results of conformationally sensitive antibodies TNT1 (B), TOC1 (C), and T22 (D). Y-axis represents % light absorbed value (converted from A450 reading). Error bars represent SD of 3 independent experiments and data sets were compared to WT 2N4R tau using an unpaired t-test. “ns” indicates no significant difference when compared to WT 2N4R control. Statistical significance was defined as *ns p* > 0.05, ∗ *p* ≤ 0.05; ∗∗ *p* ≤ 0.01; ∗∗∗ *p* ≤ 0.001.

### ARA Aggregation Kinetics

In addition to studying total aggregation, filament morphology, and immuno-reactivity, we also compared the kinetics of aggregation of each mutant to WT tau in the presence of ARA. When comparing WT tau to P270S, P332S, and P364S there was no significant difference in the maximum polymerization, rate of aggregation, and lag time (figure 3). However, the P301S mutation caused a significant increase in maximum polymerization consistant with the results of the LLS endpoint reactions (figure 3B). Although the P301S rate of polymerization was less than WT, it did not meet the statistical threshold to be considered signficantly different (figure 3C). However, P301S did cause a significant increase in the lag time when compared to WT (figure 3D). Typically, a longer lag time and slower rate of polymerization is indicative of fewer filaments with a longer average length. Therefore, the results from these aggregation kinetic experiments support, at least in part, the findings from the TEM studies (figure 1).

**Figure 3:**
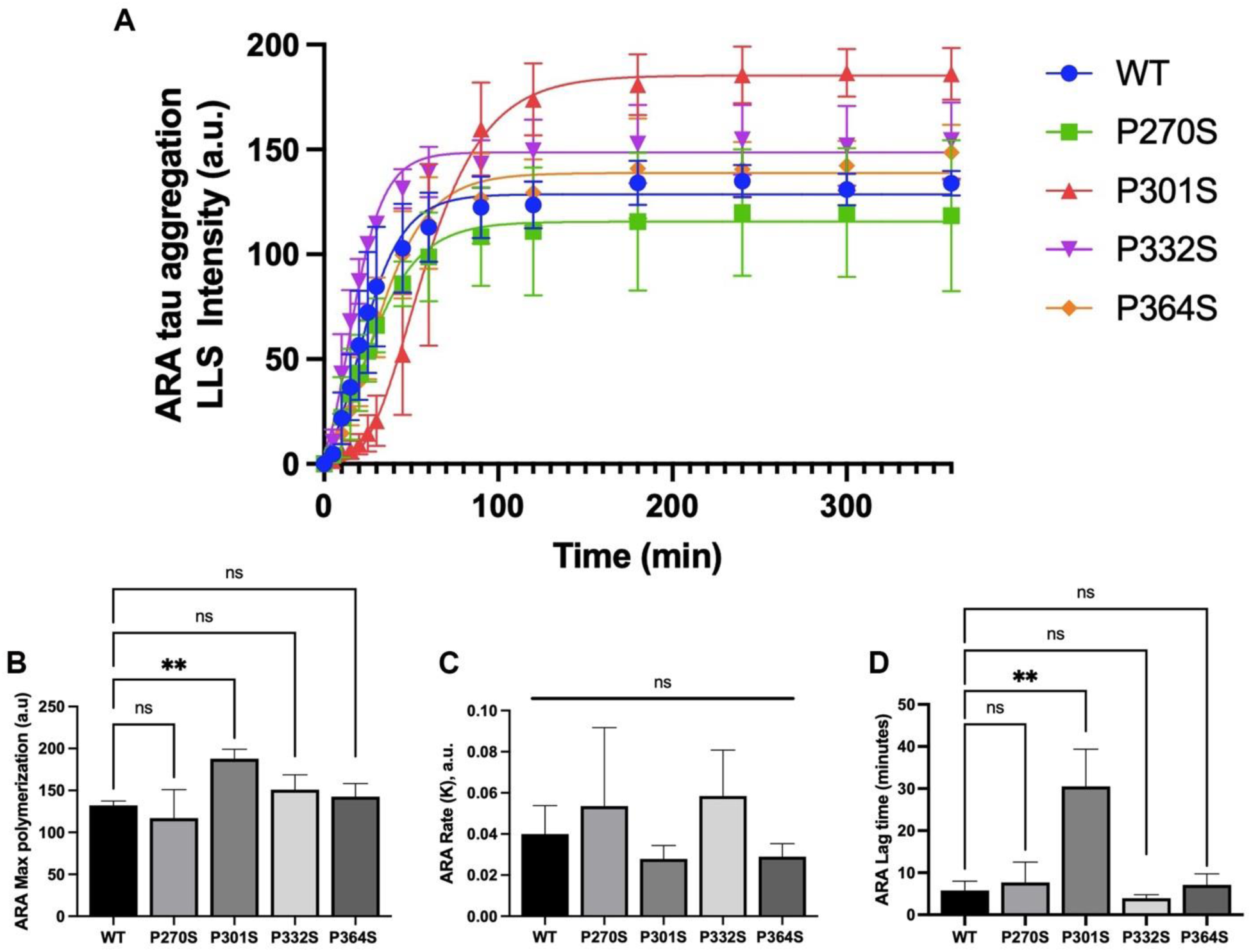
ARA induced aggregation kinetics. Three independent data sets of ARA induced reactions monitored by LLS were fit to a plateau followed by one phase association non-linear curve. A) average LLS intensity over the course of 6 hours (360 minutes) were plotted for WT (blue dots), P270S (green squares), P301S (red triangles), P332S (purple inverted triangles), and P364S (orange diamonds). B) Average maximum polymerization, C) average rate of polymerization, and D) average lag time of each mutation was shown in a bar graph (n=3 ± s.d.). ARA induced kinetic experiments were completed by Kelsey M. Hillyer. Each P to S mutation was compared to WT using an unpaired t-test. Significance was identified as *ns p* > 0.05, ∗ *p* ≤ 0.05; ∗∗ *p* ≤ 0.01; ∗∗∗ *p* ≤ 0.001. *ns* = not significant.

### ARA Results Summary

In summary, when compared to WT tau, the P270S mutation effects average filament length and average number of filaments. In contrast, P301S causes difference in total polymerization, filament length, number of filaments, T22 immuno-reactivity, and aggregation kinetics. Compared to WT, P332S also has a significant effect on total polymerization, number of filaments, and T22 reactivity. The P364S mutation had no effect on total polymerization, filament morphology as shown by TEM, and aggregation kinetics. However, there was a significant decrease in T22 reactivity.

### P100 induced endpoint aggregation experiments

In the presence of P100 as an inducer molecule, the P270S mutation resulted in a significant decrease in LLS when compared to WT (figure 4A). Also, in contrast to the ARA induced reactions, the P270S filaments were significantly longer than WT (figure 4B). Interestingly, P100 induction of P301S did not show a significant increase in aggregation as detected by both LLS and TEM (figure 4A & 4C). However, there was a decrease in the number of filaments (figure 4D). Both P332S and P364S mutations resulted in filaments that were significantly longer than WT (figure 4B). In addition, there was a significant reduction in the number of filaments and an overall decrease in filament mass as determined by TEM (figure 4C-D). However, LLS did not reveal a significant difference between either P332S or P364S when compared to WT (figure 4A).

**Figure 4:**
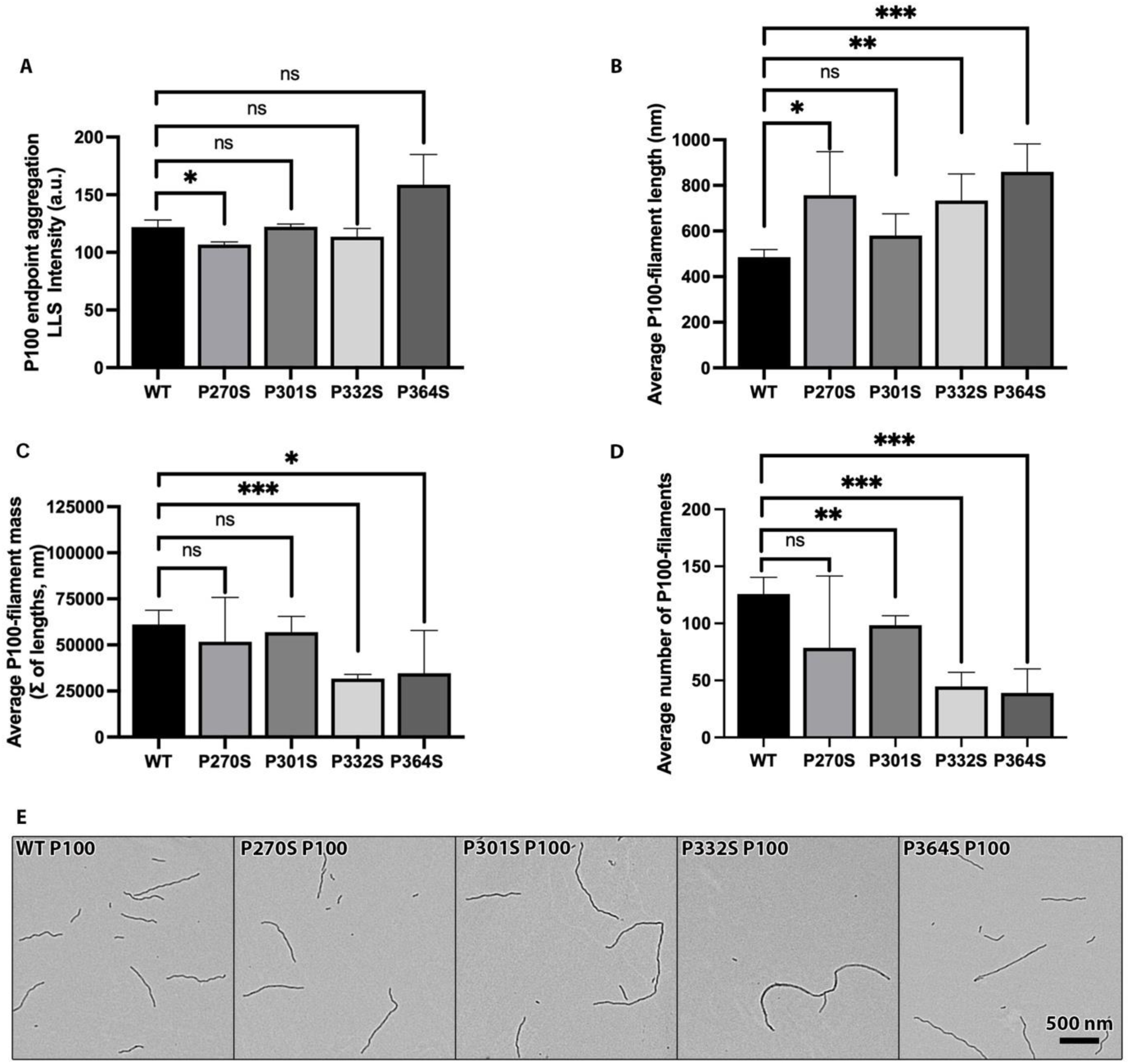
LLS and TEM endpoint measurements of P100 polyphosphate reactions. A) Endpoint total amount of aggregation by LLS (n=3 ± s.d.). 5 different random micrographs taken from a single EM grid at a 5,000x magnification were quantified to obtain measure B) average filament length (± s.d), C) average filament mass (± s.d), and D) average number of filaments (± s.d). E) Representative TEM micrographs of WT and each of the four P to S mutations (see labeling in upper left corner of each micrograph). Scale bar in bottom right corner of P364S P100 image represents 500 nm for each micrograph. Each P to S mutation was compared to WT using an unpaired t-test. *ns p* > 0.05, ∗ *p* ≤ 0.05; ∗∗ *p* ≤ 0.01; ∗∗∗ *p* ≤ 0.001. *ns* = not significant.

### P700 induced endpoint aggregation experiments

Using P700 as an inducer molecule resulted in filaments that were shorter on average than those induced with P100, and therefore more similar in average filament length to those induced by ARA (compare figure 5 with figures 1 & 4). Although, the P270S mutation did not cause any difference in the amount of LLS (figure 5A), it did result in significantly fewer, but longer, filaments when compared to WT (figures 5B &5D). In contrast to the LLS results, P270S caused a significant decrease in filament mass (compare Figure 5A & 5C). When comparing P301S induced by P700 to WT tau, there is a significant increase in LLS (figure 5A), as well as significantly fewer (figure 5D), but longer (figure 5B) filaments. These results were consistent with the ARA induced experiments, however, P301S filament mass was not statistically different than WT (figure 5C). LLS results of P332S showed no significant difference when compared to WT (figure 5A), however, TEM revealed fewer P332S filaments (figure 5D) with a much longer average filament length (figure 5B) and overall less filament mass (figure 5C). Both LLS and TEM-filament mass showed no significant difference between P364S and WT (figure 5C). Although TEM did reveal the P364S mutation to form fewer, but longer filaments when compared to WT (figure 5B & 5D).

**Figure 5:**
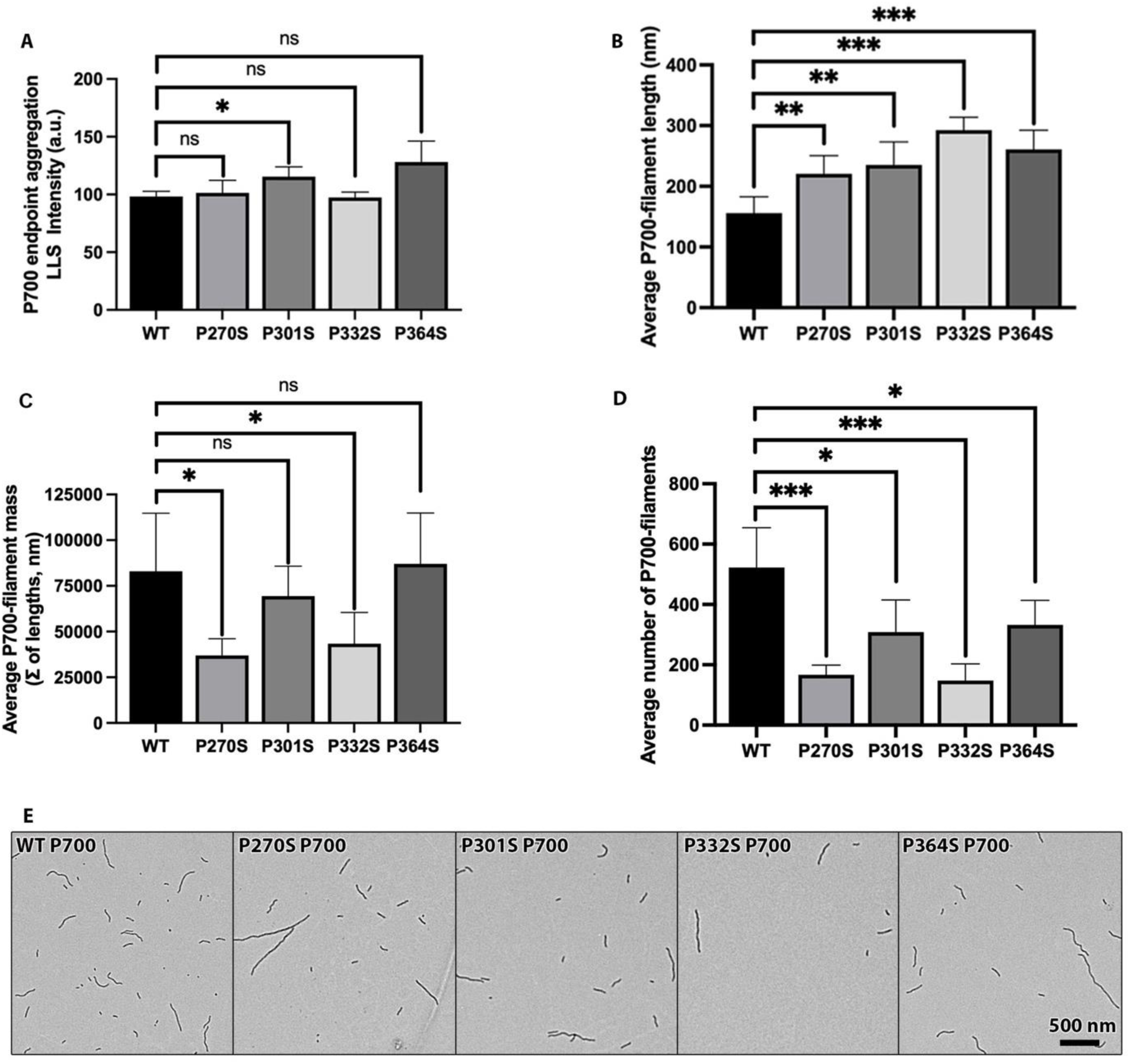
LLS and TEM endpoint measurements of P700 polyphosphate reactions. A) Endpoint total amount of aggregation by LLS (n=3 ± s.d.). 5 different random micrographs taken from a single EM grid at a 5,000x magnification were quantified to obtain measure B) average filament length (± s.d), C) average filament mass (± s.d), and D) average number of filaments (± s.d). E) Representative TEM micrographs of WT and each of the four P to S mutations (see labeling in upper left corner of each micrograph). Scale bar in bottom right corner of P364S P700 image represents 500 nm for each micrograph. Each P to S mutation was compared to WT using an unpaired t-test. *ns p* > 0.05, ∗ *p* ≤ 0.05; ∗∗ *p* ≤ 0.01; ∗∗∗ *p* ≤ 0.001. *ns* = not significant.

### Polyphosphate induced aggregation measured by sandwich ELISA

Similar to filaments induced by ARA, we saw no significant difference between WT and any of the P to S mutations using the 5,7,12 capture antibody ELISA when induced with either P100 or P700 (see figure S3). In addition, we also saw no significant effect from any of the mutants in either the TNT1 and TOC1 ELISAs using both polyphosphate inducers (figure 6). However, it does appear that the TOC1 antibody has a lower affinity towards polyphosphate induced filaments than ARA induced filaments (see figure S4 comparing WT aggregates in the presence of each inducer). However, the most striking difference in the immuno-reactivity experiments was seen using T22 as a capture antibody. Both WT (figure S4) and each of the P to S mutants (data not shown) induced by either P100 or P700 had no reactivity towards T22 in the ELISA experiment. Therefore, we were unable to compare the effect of the mutations using this particular antibody. We suspected that this may be due to the inducer molecule blocking the T22 binding site. This was tested by performing a dot-blot experiment to compare both WT and P301S induced by ARA, P100, P700, long RNA, and small RNA and detect using T22. In this experiment, T22 did detect the polyphosphate and RNA induced aggregates (figure S5).

**Figure 6:**
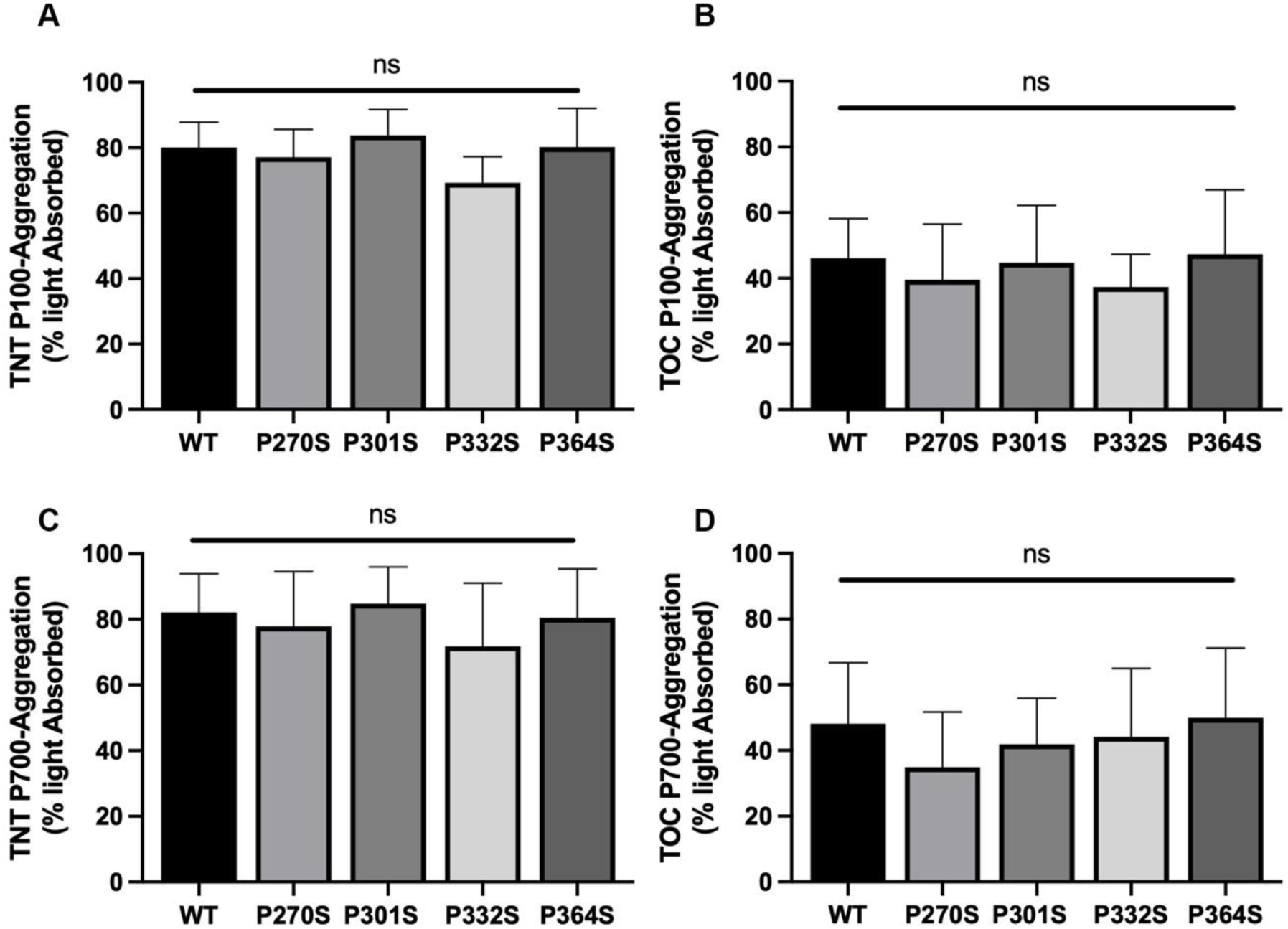
Polyphosphate TNT and TOC ELISA data. ELISA results using TNT (A & C) and TOC (B & D) capture antibodies against P100 (A & B) and P700 (C & D) induced filaments. Y-axis represents % light absorbed value (converted from A450 reading). Error bars represent SD of 3 independent experiments. Each P to S mutation was compared to WT using an unpaired t-test. *ns p* > 0.05, ∗ *p* ≤ 0.05; ∗∗ *p* ≤ 0.01; ∗∗∗ *p* ≤ 0.001. *ns* = not significant.

### P100 aggregation kinetics

Similar to the ARA aggregation studies, we used LLS to monitor protein aggregation over time. Compared to WT tau, P332S had a significant decrease in rate of polymerization (*K*), but was similar in terms of maximum polymerization (figure 7). Although the difference in lag time between WT and both P270S and P332S was not statistically significant due to the high variability in lag time, the average was much higher than WT, which had essentially no lag period (figure 7D). Similarly, P301S also appeared to have no lag period and total polymerization was similar to WT (figure 7B & 7D). However, the rate of polymerization for P301S was much faster than WT (figure 7C). This is in stark contrast to the results seen from the ARA induced aggregation kinetic experiments, which showed P301S to have a longer lag time, greater maximum polymerization, and no significant difference in rate (Compare figure 3 to figure 7). The P364S mutation caused a significant decrease in maximum polymerization and a significant increase in the lag time, but no change to rate of polymerization (figure 7).

**Figure 7:**
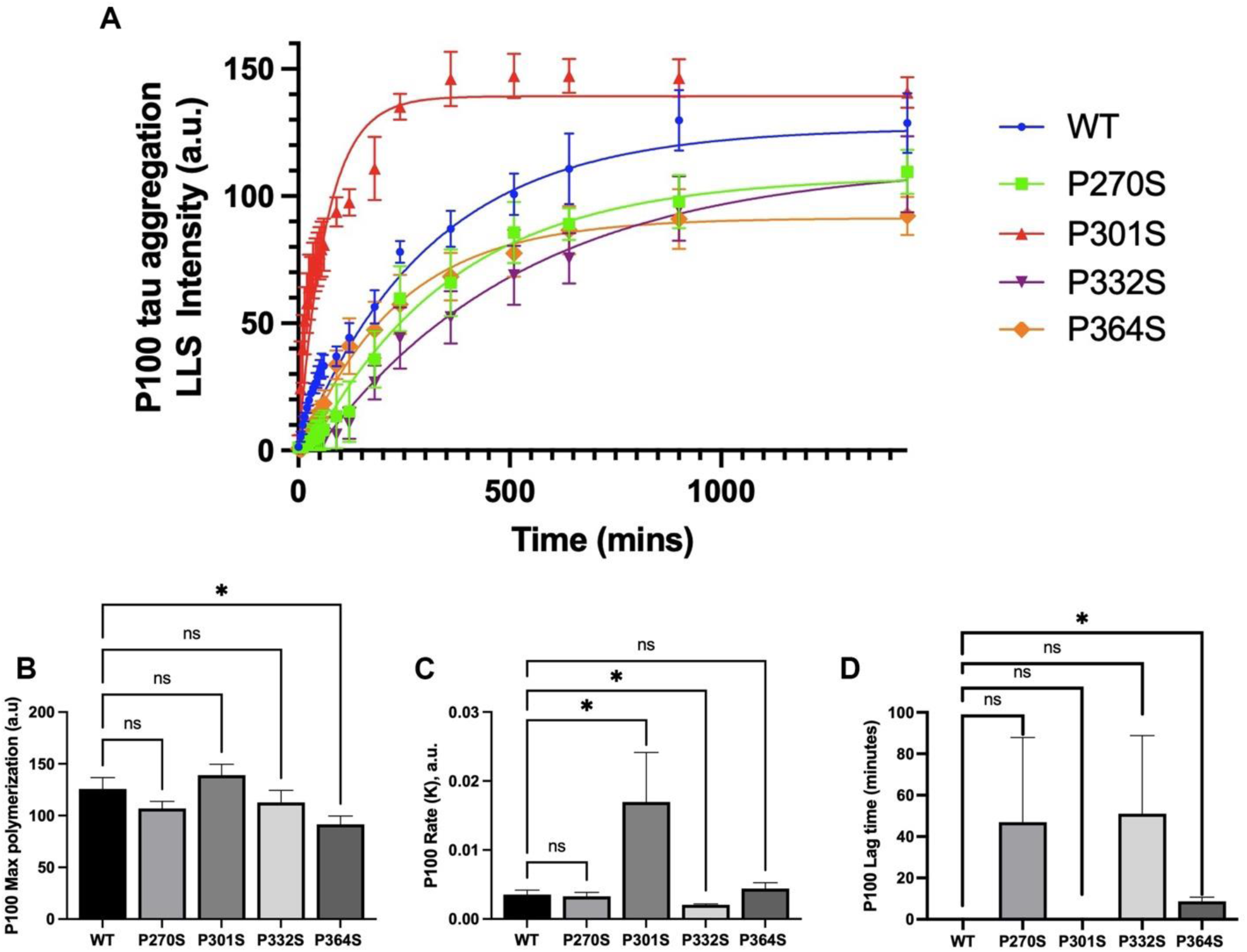
P100 induced aggregation kinetics. Three independent data sets of P100 induced reactions monitored by LLS were fit to a plateau followed by one phase association non-linear curve. A) average LLS intensity over the course of 24 hours (1440 minutes) were plotted for WT (blue dots), P270S (green squares), P301S (red triangles), P332S (purple inverted triangles), and P364S (orange diamonds). B) Average maximum polymerization, C) average rate of polymerization, and D) average lag time of each mutation was shown in a bar graph with error bars representing standard deviation. Each P to S mutation was compared to WT using an unpaired t-test. *ns p* > 0.05, ∗ *p* ≤ 0.05; ∗∗ *p* ≤ 0.01; ∗∗∗ *p* ≤ 0.001. *ns* = not significant.

### P700 aggregation kinetics

In the presence of P700 as an inducer molecule, the P270S mutation did not significantly affect maximum polymerization (figure 8). P270S did have a significantly faster rate of polymerization when compared to WT (figure 8C), and although not significant, the average lag time was higher than WT (figure 8D). In the case of P301S, the P700 inducer had the same effect as P100 as shown by the significant increase in the rate of polymerization (figure 8C), but no change in maximum polymerization or lag time (figure 8B & 8D). In contrast to the P100 induced kinetic experiments, P332S was the only mutation that caused a significant decrease in maximum polymerization (figure 8B). This difference was particularly striking as it was approximately 1/3 of the WT maximum polymerization. Despite this large difference, there was no change in the rate of polymerization, although the increase in lag time was statistically significant (figure 8C & 8D). In contrast to the results from the P100 induced experiments, but more similar to the ARA induced experiments, P364S had no significant effect on maximum polymerization, rate, or lag time (figure 8).

**Figure 8:**
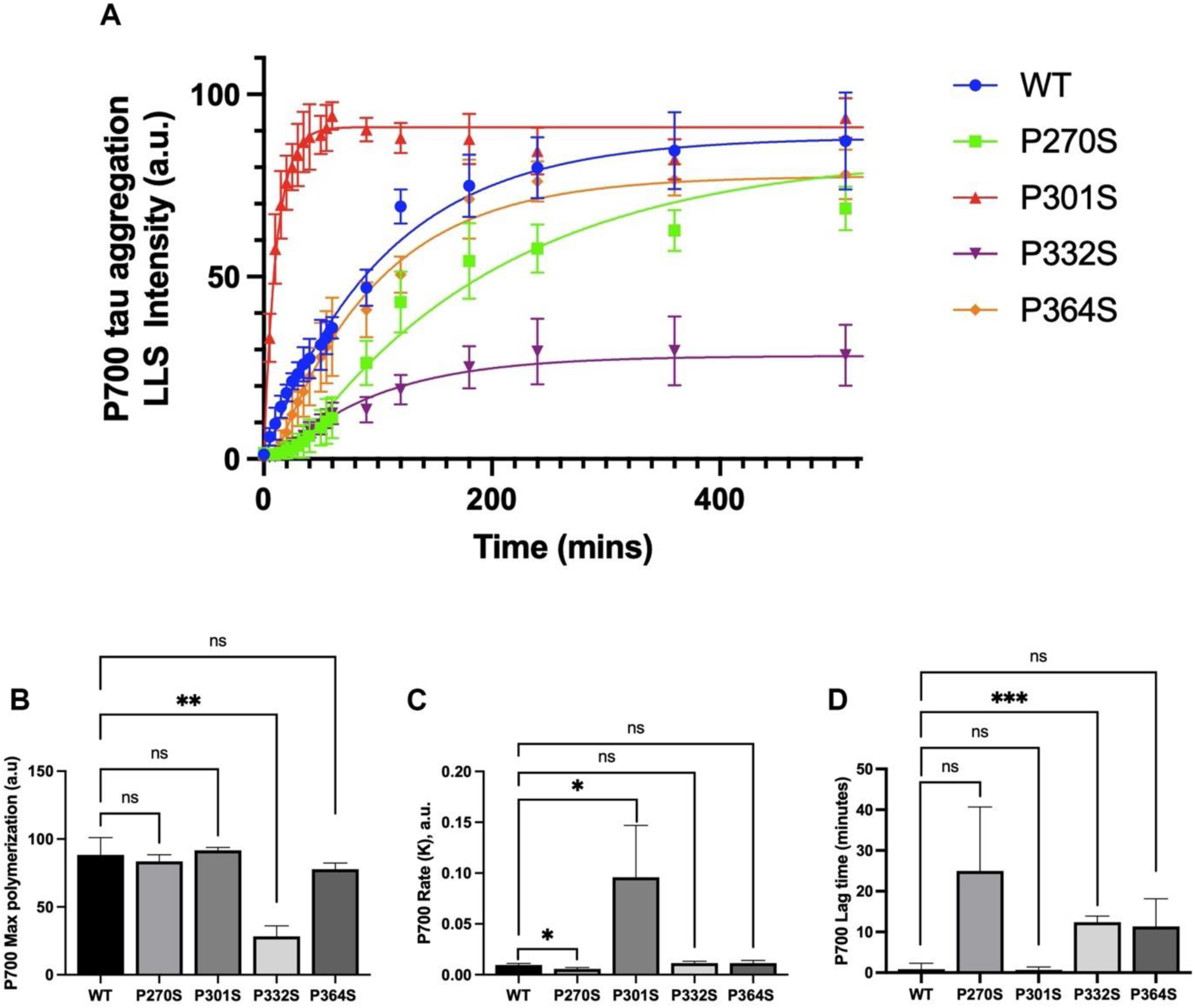
P700 induced aggregation kinetics. Three independent data sets of P700 induced reactions monitored by LLS were fit to a plateau followed by one phase association non-linear curve. A) average LLS intensity over the course of 8.5 hours (510 minutes) were plotted for WT (blue dots), P270S (green squares), P301S (red triangles), P332S (purple inverted triangles), and P364S (orange diamonds). B) Average maximum polymerization, C) average rate of polymerization, and D) average lag time of each mutation was shown in a bar graph with error bars representing standard deviation. Each P to S mutation was compared to WT using an un-paired t-test. Significance was identified as *ns p* > 0.05, ∗ *p* ≤ 0.05; ∗ ∗ *p* ≤ 0.01; ∗∗∗ *p* ≤ 0.001. *ns* = not significant.

### Polyphosphate Summary

The P270S mutation, previously shown to have no effect on total aggregation, a decrease of average filament length, and an increase in the number of filaments when induced by ARA, causes a decrease in total aggregation as measured by LLS and an increase in average filament length as measured by TEM when induced with P100. When P270S is induced with P700, the average filament length is still longer, but there are fewer filaments and less overall filament mass. In addition, analysis of P700 induced kinetic data revealed a decrease in rate of polymerization for P270S. The P301S mutation causes a decrease in the number of filaments as measured by TEM and a significant increase in rate of polymerization when induced with both P100 and P700. When induced with P700, there was an increase in total aggregation as determined by LLS, and an increase in average filament length as measured by TEM. Both P100 and P700 induced P332S reactions showed similar changes in filament length, mass, and number of filaments. P100 induced P332S caused a decrease in rate of polymerization and P700 induced filaments showed a decrease in the maximum polymerization and increase in lag time. While both P100 and P700 induced P364S reactions showed an increase in average filament length and decrease in the number of filaments, only P100 induced reactions had a negative effect on total filament mass and maximum polymerization (as determined by kinetic analysis). In addition, P100 induced P364S also caused a decrease in maximum polymerization and increase in lag time as shown by kinetic analysis. None of these differences were shown when inducing P364S with ARA. Both WT and each of the P to S mutations induced with either P100 or P700, did not react with T22 in the ELISA experiment.

### RNA induced aggregation

Endpoint aggregation experiments were performed using small RNA and long RNA as inducers and the analysis of aggregation was completed using LLS and TEM (figure 9 and figure 10) similar to ARA, P100, and P700. WT tau and many of the mutants had total amounts of aggregation that were much lower than reactions induced with ARA, P100, and P700 (compare to figures 1, 4, & 5).

**Figure 9:**
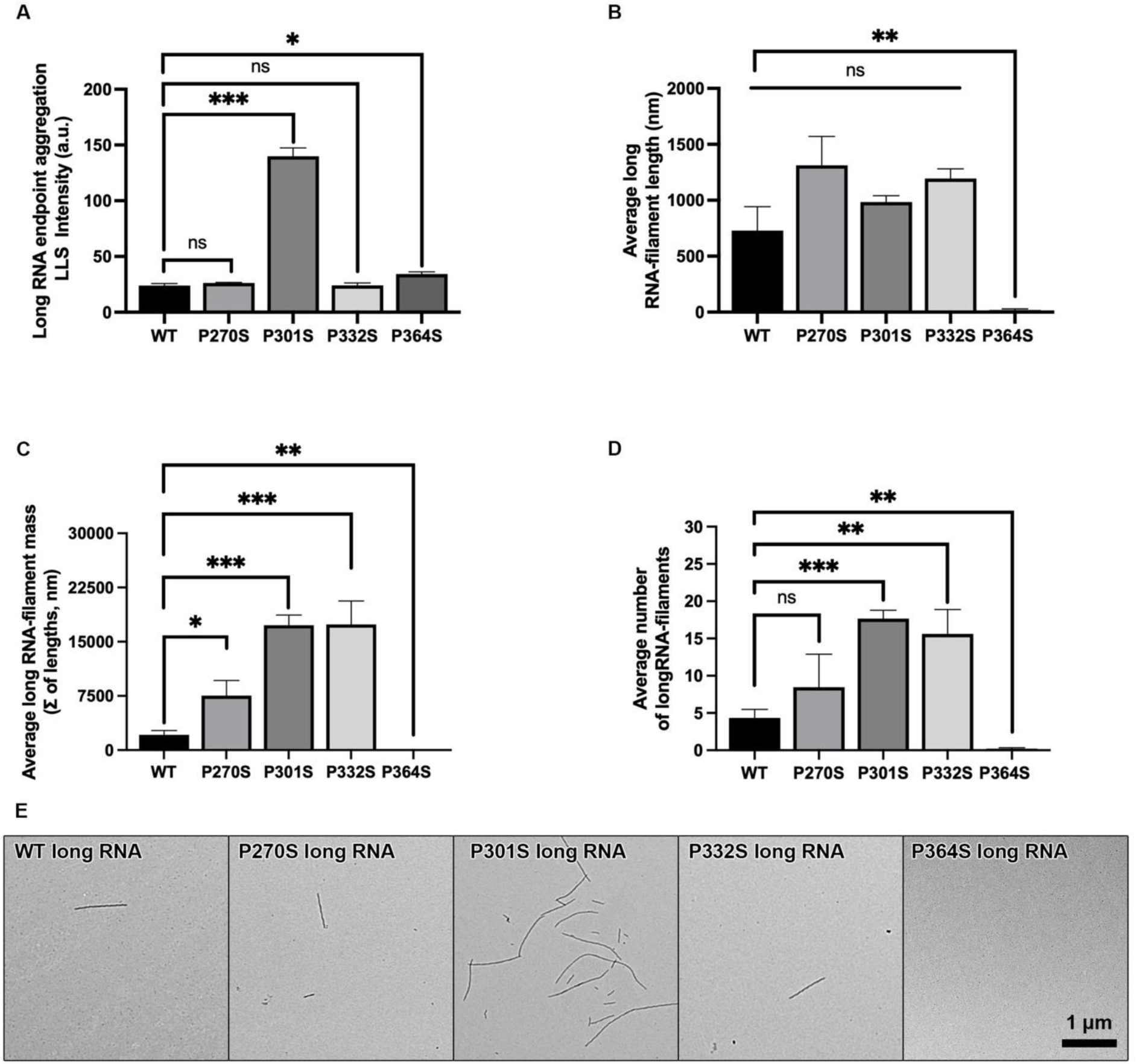
LLS and TEM measurements of long RNA endpoint aggregation reactions. LLS measurement of endpoint total amount of aggregation of three independent experiments (A). B-D quantitative EM results, average filament length (B), average filament mass (C), average number of filaments (D). E) Representative TEM micrographs of WT and each of the four P to S mutations (see labeling in upper left corner of each micrograph). Scale bar in bottom right corner of P364S long RNA image represents 1 µm for each micrograph. Figures B-D represent averages of 15 different random micrographs taken from a single EM grid at a 5,000x magnification. LLS and TEM grid preparation was completed by Madison J. McGuire. Error bars on figures A-D represents SD of each data set. Each P to S mutation was compared to WT using an unpaired t-test. ∗ *p* ≤ 0.05; ∗∗ *p* ≤ 0.01; ∗∗∗ *p* ≤ 0.001.

**Figure 10:**
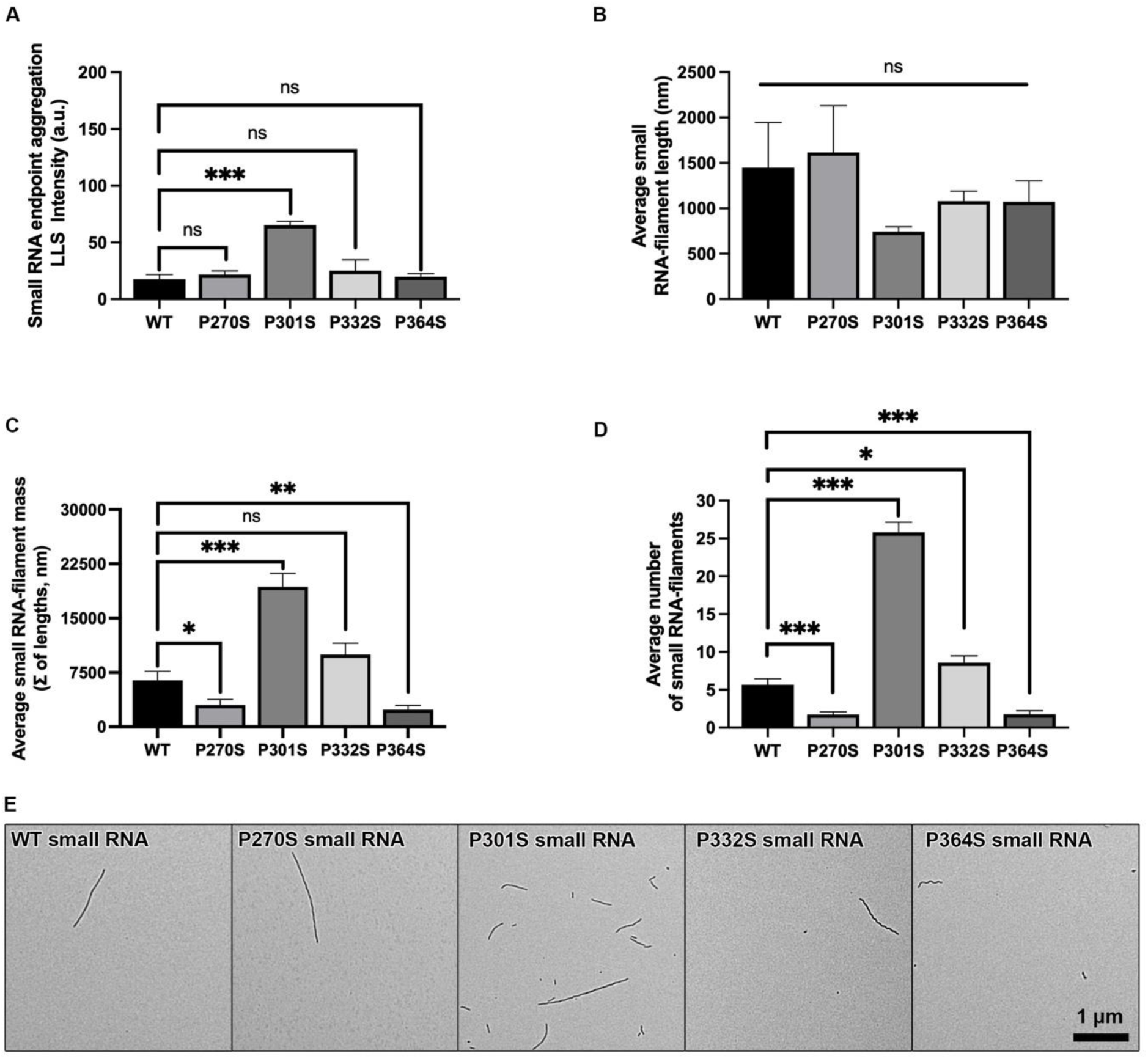
LLS and TEM measurement of small RNA induced endpoint aggregation reactions. LLS measurement of endpoint total amount of aggregation of three independent experiments (A). B-D quantitative EM results, average filament length (B), average filament mass (C), average number of filaments (D). E) Representative TEM micrographs of WT and each of the four P to S mutations (see labeling in upper left corner of each micrograph). Scale bar in bottom right corner of P364S small RNA image represents 1 µm for each micrograph. Figures B-D represent averages of 15 different random micrographs taken from a single EM grid at a 5,000x magnification. Error bars on figures A-D represents SD of each data set. Each P to S mutation was compared to WT using an unpaired t-test. ∗ *p* ≤ 0.05; ∗∗ *p* ≤ 0.01; ∗∗∗ *p* ≤ 0.001.

### Long RNA endpoint aggregation experiments

In contrast to the results using ARA, P100, and P700, total filament mass as determined by TEM (figure 9C) showed that the P270S mutation caused a significant increase in aggregation when induced with long RNA. As measured by LLS, the P301S mutation caused a significant increase of more than 5-fold when compared to WT tau (figure 9A). This was further supported by an approximate 5-fold increase in filament mass (figure 9C). In contrast to the results of the ARA induced reactions, P301S induced with long RNA did not result in increased filament length, but did result in an increase in the total number of filaments (compare figure 1 B & D to figure 9 B&D). Similarly, the P332S mutation also resulted in an increase in filament mass and number (figure 9 C & D). Although P364S induced reactions did result in significantly higher light scattering (figure 9A), no filaments were detected using TEM (figure 9 B-E). This suggests that either long RNA induced P364S aggregates are not stable and depolymerize during TEM grid preparation, filaments are below the TEM detection threshold (<25 nm), that RNA interacts with P364S causing it to scatter light, but not form filaments, or that long RNA induced P364S aggregates have properties that reduce their adherence to EM grids.

### Small RNA endpoint aggregation experiments

Although endpoint aggregation measured by LLS (figure 10A) showed the P270S mutation had no effect on aggregation when compared to WT tau, TEM analysis revealed a significant decrease in filament mass and number of filaments (figure 10 C & D, respectively). When comparing small RNA induced P301S and WT tau, the results were similar to the results when inducing with long RNA. However, the increase in aggregation caused by the P301S mutation was not as stark using small RNA (∼3 fold) when compared to reactions induced with long RNA (>5 fold). Using small RNA as an inducer, the P332S mutation resulted in no significant difference in LLS, length of filaments, and filament mass, however there was a significant increase in the number filaments (figure 10 A-D). In contrast to the TEM results of P364S induced with long RNA, using small RNA as an inducer did form filaments that were detected using TEM (figure 10 B-E). However, total filament mass and number of filaments formed was significantly less than WT tau (figure 10 C & D).

### RNA induced aggregation measured by sandwich ELISA

Despite a noticeable difference between P301S and WT tau as shown by TNT ELISA for long RNA, due to the variability in data this result did not reach statistical significance (figure 11). This was also the case for both long RNA and small RNA when comparing TOC1 ELISA results. However, small RNA induced P301S TNT1 reactivity was significantly higher than WT tau. None of the other mutations were significantly different than WT with regard to either the TNT1 and TOC1 ELISA using both long RNA and small RNA.

**Figure 11:**
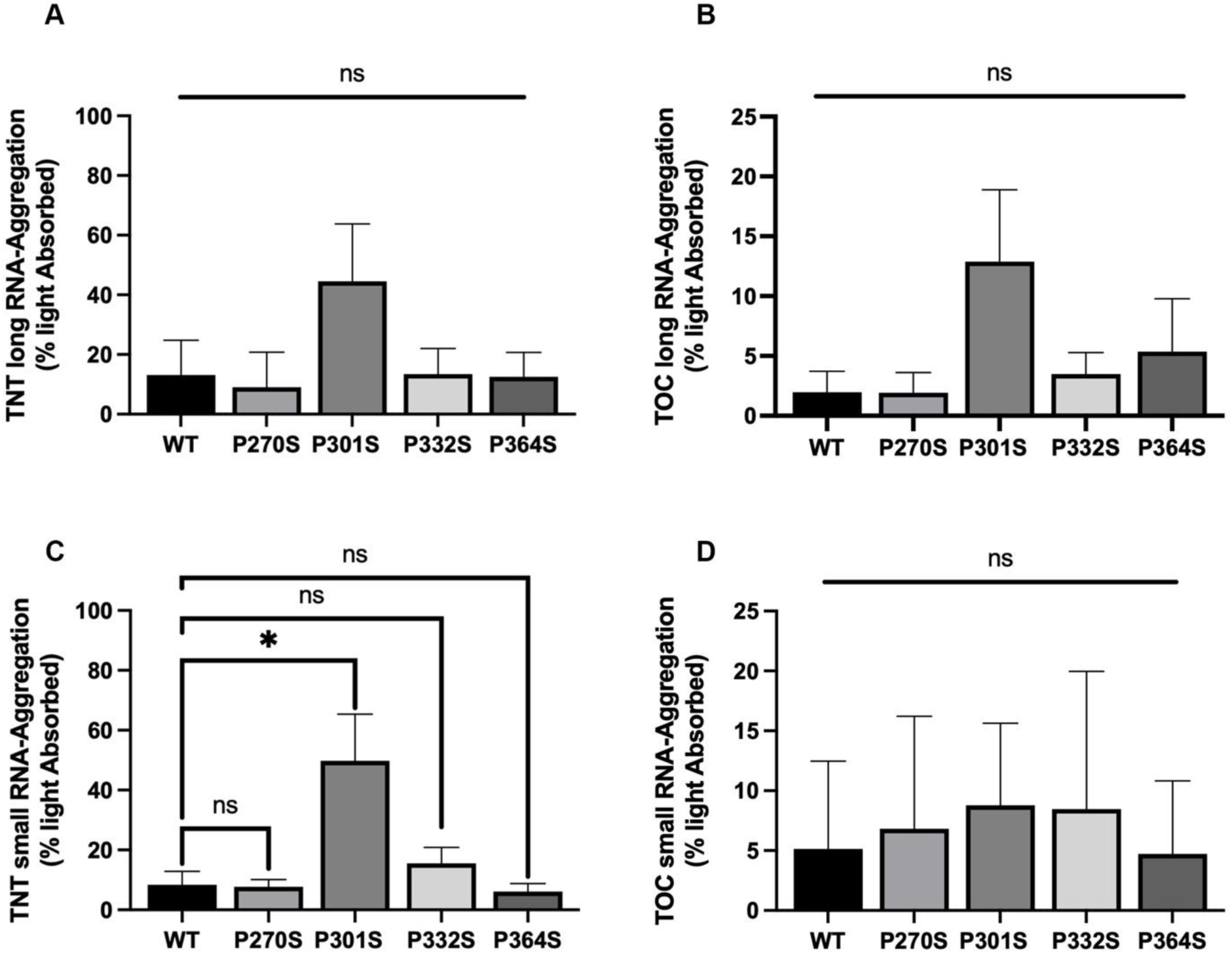
Immuno-reactivity of long RNA and small RNA reactions. ELISA results using TNT (A & C) and TOC (B & D) capture antibodies against long RNA (A & B) and small RNA (C & D) induced filaments. Y-axis represents % light absorbed value (converted from A450 reading). Error bars represent SD of 3 independent experiments. Each P to S mutation was compared to WT using an unpaired t-test. *ns p* > 0.05, ∗ *p* ≤ 0.05; ∗∗ *p* ≤ 0.01; ∗∗∗ *p* ≤ 0.001. *ns* = not significant.

### RNA summary

Using both long RNA and small RNA as an inducer molecule, we found differences in total polymerization, filament length, filament mass, number of filaments, and immuno-reactivity. The P270S mutation caused no change in total polymerization as determined by LLS for both long RNA and small RNA. However, when induced with small RNA there was a significant decrease in filament mass and number of filaments as determined by TEM. In contrast, P270S induced with long RNA caused a significant increase in filament mass, and an increase in the number of filaments and average filament length, although neither were statistically significant. There was also no significant difference in immuno-reactivity between P270S and WT using either TOC1 or TNT1 capture antibodies for either sizes of RNA.

When inducing P301S with both small and long RNA, there was a dramatic increase in total aggregation as determined by LLS (>5 fold for long RNA, and ∼3 fold for small RNA). Interestingly, despite an increase in LLS, number of filaments, and total filament mass, using both RNA inducer molecules, the P301S mutation did not cause a significant difference in average filament length when compared to WT. Using long RNA to induce P301S aggregation caused an increase in TNT1 and TOC1 reactivity, however this increase did not reach the threshold to be statistically significant. Using small RNA, there was a significant increase in TNT1 reactive species, however no substantial change was seen when using the TOC1 capture antibody. The P332S mutation had no significant effect on total polymerization as determined by LLS, however, TEM analysis revealed an increase in number of filaments when induced with both RNA species, and a significant increase in filament mass when induced with long RNA. In initial endpoint aggregation experiments, there was a significant increase in LLS when induced with long RNA, however, no filaments were present in the EM micrographs. When comparing P364S and WT induced with small RNA, there was no significant change in LLS and filaments were present as shown by TEM. Analysis of these images revealed a decrease in total filament mass and number of filaments, but no significant change to average filament length.

### Comparison of each inducer molecule

Table 1 summarizes the results of each experiment using each of the 5 inducer molecules: ARA, P100, P700, long RNA, and small RNA). It is clear from this summary that the choice of inducer molecule and method of aggregate detection can influence whether differences are detected and also the absolute extent of differences. It is also apparent that the P301S more consistently demonstrates differences from the wild type protein regardless of inducer and method of detection as compared to the other P to S mutations.

**Table 1:** Comparison of *in vitro* aggregation results by inducer, mutant, and method of detection.

| Method<br>Parameter | LLS | TEM |  |  | sELISA |  |  | Kinetics |  |  |
| --- | --- | --- | --- | --- | --- | --- | --- | --- | --- | --- |
| | Max | $\bar{L}$ | $\Sigma$ | # | TNT1 | TOC1 | T22 | Max | Rate | Lag |
| *Arachidonic Acid* |  |  |  |  |  |  |  |  |  |  |
| P270S | ≈ | --- | ≈ | + | ≈ | ≈ | ≈ | ≈ | ≈ | ≈ |
| P301S | ++ | ++ | ≈ | --- | ≈ | ≈ | --- | ++ | ≈ | ++ |
| P332S | - | ≈ | --- | -- | ≈ | ≈ | --- | ≈ | ≈ | ≈ |
| P364S | ≈ | ≈ | ≈ | ≈ | ≈ | ≈ | - | ≈ | ≈ | ≈ |
| *Polyphosphate P100* |  |  |  |  |  |  |  |  |  |  |
| P270S | - | + | ≈ | ≈ | ≈ | ≈ | n.a. | ≈ | ≈ | ≈ |
| P301S | ≈ | ≈ | ≈ | -- | ≈ | ≈ | n.a. | ≈ | + | ≈ |
| P332S | ≈ | ++ | --- | --- | ≈ | ≈ | n.a. | ≈ | - | ≈ |
| P364S | ≈ | +++ | - | --- | ≈ | ≈ | n.a. | - | ≈ | + |
| *Polyphosphate P700* |  |  |  |  |  |  |  |  |  |  |
| P270S | ≈ | ++ | - | --- | ≈ | ≈ | n.a. | ≈ | - | ≈ |
| P301S | + | ++ | ≈ | - | ≈ | ≈ | n.a. | ≈ | + | ≈ |
| P332S | ≈ | +++ | - | --- | ≈ | ≈ | n.a. | -- | ≈ | +++ |
| P364S | ≈ | +++ | ≈ | - | ≈ | ≈ | n.a. | ≈ | ≈ | ≈ |
| *Long RNA (> 200 nts)* |  |  |  |  |  |  |  |  |  |  |
| P270S | ≈ | ≈ | + | ≈ | ≈ | ≈ | n.a. | n.a. | n.a. | n.a. |
| P301S | +++ | ≈ | +++ | +++ | ≈ | ≈ | n.a. | n.a. | n.a. | n.a. |
| P332S | ≈ | ≈ | +++ | ++ | ≈ | ≈ | n.a. | n.a. | n.a. | n.a. |
| P364S | + | --- | -- | -- | ≈ | ≈ | n.a. | n.a. | n.a. | n.a. |
| *Small RNA (<200 nts)* |  |  |  |  |  |  |  |  |  |  |
| P270S | ≈ | ≈ | - | --- | ≈ | ≈ | n.a. | n.a. | n.a. | n.a. |
| P301S | +++ | ≈ | +++ | +++ | + | ≈ | n.a. | n.a. | n.a. | n.a. |
| P332S | ≈ | ≈ | ≈ | + | ≈ | ≈ | n.a. | n.a. | n.a. | n.a. |
| P364S | ≈ | ≈ | -- | --- | ≈ | ≈ | n.a. | n.a. | n.a. | n.a. |

This is a summary of the results from inducer aggregation experiments. $\bar{L}$ is average filament length; $\Sigma$ is total filament mass; # is the number of filaments per micrograph; ≈ is no significant change; + indicates $p \leq 0.05$ significant increase from wt; ++ is $p \leq 0.01$ ; and +++ is $p \leq 0.001$ ; - indicates $p \leq 0.05$ significant decrease from wt; -- is $p \leq 0.01$ ; and --- is $p \leq 0.001$ ; n.a. (not applicable) indicates that the method could not be used for those conditions.

## Discussion

It has been known for several decades that the term tauopathy includes a wide range of neurological disorders with diverse etiology, clinical presentation, and histopathology. Recent advances in structural biology techniques, primarily cryo-electron microscopy, have now shown that different tauopathies also include a range of structurally diverse tau aggregates^12, 13, 34, 35^.

Previous studies have shown that missense mutations of the *MAPT* gene are associated with effects on tau pathology, neurodegeneration, and neurotoxicity^36–38^. Here, we examine how the use of different inducer molecules can have a profound effect on our understanding of the molecular characteristics of wild type tau and how it relates to missense mutations associated with a group of familial tauopathies known as frontotemporal dementia with parkinsonism linked to chromosome-17 (FTDP-17).

Previously, we, and others, have completed extensive *in vitro* aggregation studies using ARA as an inducer molecule of recombinantly expressed human tau. This has allowed for the development of optimized aggregation conditions and standard techniques to be able to complete characterization studies of *MAPT* mutations and other protein modifications. Comparisons of different fatty acids to induce tau aggregation have also shown that alkyl chain length and saturation of the fatty acid used can influence filament length, rate of polymerization, amount of polymerization, and filament density^39, 40^. In this study, we have used ARA as a fatty acid inducer of tau aggregation as it forms filaments that have similar morphology to straight filaments isolated from AD and has relatively higher stability when compared to fatty acids with a less saturated aliphatic tail^16^.

Following ARA induced aggregation experiments, we sought to compare the P to S mutations using polyphosphate as an inducer molecule. Similar to fatty acids, chain lengths of polyphosphate can influence the amount and length of tau filaments^22^. However, there are fewer studies characterizing these differences. For the purpose of this study we chose to complete aggregation experiments using medium chain polyphosphate (P100) and long chain polyphosphate (P700). Similarly, few studies have characterized how RNA induces tau aggregation. To limit the scope of this study we chose to use two different RNA samples that were separated based on length. However, it is possible that differences in small and long RNA’s primary and secondary structure may be the cause of the variance of observed effects on aggregation.

In this study we used three independent assays (LLS, TEM, and ELISA) for studying the effects of disease related mutations in the PGGG motif in order to understand how these effects relate to the inducer molecule used. Right-angle laser light scattering (LLS) is a standard tool used in the field of studying *in vitro* protein aggregation. However, this assay does have certain limitations when being used to compare different samples. For example; 1) samples with dissimilar size distributions will scatter light dis-proportionately to each other, 2) changes in filament flexibility could potentially affect the radius of gyration and therefore the total amount of light scattered, 3) formation of micelles or RNA droplets could interfere with measuring filament light scattering, and 4) particles less than 1/20^th^ the wavelength of the light source will not scatter light proportionately. In addition, when light scattering endpoint aggregation studies are completed the reactions take place in a closed plastic microcentrifuge tube, but when light scattering is used for measuring aggregation kinetics, the reactions are incubated in a 5 mm x 5 mm glass cuvette sealed with parafilm. The differences in the material of these vessels and how each one affects surface area to volume ratio, may cause significant effects on total aggregation. For example, the P332S mutation when induced with P700 has no significant effect on total aggregation as measured by LLS during the endpoint aggregation reactions. However, during aggregation kinetic studies, the maximum polymerization of P332S was significantly reduced when compared to WT tau.

In addition to using electron microscopy to validate the results of the LLS aggregation assay, we were also interested in finding out more information such as the total filament mass, average length of filaments, and number of filaments using each inducer. Through the use of image analysis software, we are able to use transmission electron microscopy as a semi-quantitative tool, with the following limitations. 1) Only filaments that bind to the EM grid are detectable and it is possible that certain tau variants and/or inducer-specific polymorphs will have a different binding affinity toward the EM grid. 2) Filament length may be altered by filament breaking as they adhere to the grid. 3) The stain used to visualize the filaments may bind to different tau variants and/or inducer-specific polymorphs differently. 4) We are visualizing a relatively small sample size of filaments compared to the sample of those analyzed by LLS^41^.

Sandwich ELISA assays can provide useful information about the species of tau aggregate being used and how these might relate to toxic species associated with disease. Similar to LLS and quantitation TEM, ELISA also has certain assay specific limitations when comparing multiple different samples. 1) The aggregates must form the corresponding epitope to the antibodies being used, and that epitope must be accessible, 2) inducer molecules may interfere with antibody binding by blocking epitope recognition, 3) changes in filament flexibility may change epitope accessibility, 4) changes in length distribution may change the number of epitopes accessible (this is known to be the case with TOC1, as it is thought to bind to filament ends^19^), 5) may not appear to validate other methods due to different limits of detection (oligomers smaller than 1/20^th^ LLS wavelength and 25 nm in length, may still bind to oligomer specific antibodies such as TOC1 and T22, and pro-aggregate monomer may still bind TNT1). Therefore, with these limitations in mind, it is important be cautious when basing conclusions on any single assay. In addition to actual changes to filaments caused by FTDP-17 mutations, we should also consider the disproportionate effect mutations may have on the method of detection.

Initial endpoint aggregation studies using ARA, P100, P700, long RNA, and small RNA, illustrates that the type of inducer used can have significant effects on the total amount of aggregation as well as the effect missense mutations can have on total aggregation relative to wild type tau (WT). Although the P301S mutation caused a significant increase in ARA-induced aggregation when compared to WT, there was no statistically significant difference when induced with P100 polyphosphate. Interestingly, using P100 as an inducer, the P270S mutation did result in a significant decrease in aggregation, however this result was not seen with any of the other inducers. When induced with the longer chain P700 polyphosphate, a significant difference was seen when comparing P301S to WT. Although these results were reproducible among independent experiments, they were relatively subtle with a difference of less than 20% when comparing WT to P301S (ARA and P700) or P270S (P100). In contrast, the effect of the P301S mutation on RNA induced aggregation was much more noticeable. In the case of small RNA, P301S aggregated approximately 3-fold more when compared to WT tau. Similarly, when induced with long RNA, P301S caused over a 5-fold increase in aggregation.

In addition, LLS experiments showed the P332S mutation increased aggregation when induced with ARA. This was further supported by the significant increase in filament mass as shown by TEM. Interestingly, P364S induced with long RNA also showed a significant increase in light scattering, however TEM analysis did not reveal any filaments, suggesting the possibility that this mutation enhances tau’s interaction with RNA, but not resulting in the formation of stable fibrils.

Transmission electron microscopy revealed that filaments induced with the shorter chain P100 polyphosphate had a much longer average filament length than those induced with long chain P700 polyphosphate (compare figure 5 B and E to figure 6 B and E). Further comparisons of P to S mutations on filament length show substantial differences between the different inducer classes. For example; the effect of the P364S mutation on total aggregation, average filament length, and number of filaments when induced with ARA is not significantly different from WT. However, in the case of both P100 and P700 there appears to be significantly fewer, but longer filaments.

The influence of polyphosphate chain length on average tau filament length has previously been studied from a biophysical perspective^22^. This finding is further supported by our TEM and kinetic studies. For example, in the case of P100 induced aggregates average polymerization rates of WT and each of the P to S mutants are slower than the rate of their respective P700 induced counterparts. This suggests that filaments induced with P100 form more slowly but result in longer filaments than those induced with P700.

The filaments induced with both P100 and P700 appear to follow a nucleation independent mechanism, or a fast nucleation step as shown by the extremely short lag time, especially in the case of WT and P301S tau. Although, further investigations would be required to fully elucidate the mechanism of polyphosphate aggregation. Interestingly, the P301S mutation causes an increase in maximum aggregation for ARA, a slight but not significant increase for P100, and no change to P700 induced filaments. However, in the case of ARA induced filaments P301S has a significantly longer lag time with a slightly slower rate of aggregation, when compared to WT. This increased lag time suggests the P301S mutation slows the nucleation step. Conversely, in the case of both P100 and P700, neither WT, nor P301S have a measurable lag time, but the P301S rate of aggregation is substantially faster. This shows a fundamental difference between polyphosphate and ARA induced filaments in regards to the effect of the P301S mutation on aggregation kinetics and therefore warrants further investigation.

When completing kinetic studies using long RNA (figure S6), and to a lesser extent small RNA (data not shown), as an inducer molecule we witnessed a strange phenomenon where initial addition of the inducer caused almost immediate light scattering that then faded over a period of approximately 28 hours. This was then followed by a steady increase of light scattering between 28 and 72 hours. Samples prepared for TEM imaging at time zero showed no filaments present. However, images of samples at the 72-hour time point showed a proportional amount of tau filaments to the amount of light scattering. Initial light scattering may be due to an immediate interaction between RNA and monomeric tau that is then followed by a dissociation and subsequent filament formation. Alternatively, initial light scattering could be RNA acting as a crowding agent causing liquid-liquid phase separation that form highly concentrated droplets of monomeric tau that are able to scatter light. However, more extensive studies would be required to fully understand this process and was considered outside the scope of this initial investigation.

Of the 9 tau based therapeutics that are currently in clinical trials, 6 are either passive or active immunotherapies^42^. Understanding the effect of different disease related mutations and *in vitro* inducer molecules have on antibody affinity toward tau aggregates can therefore not only provide useful characterization information, but may also be useful in the development of future therapeutics. From our current study, the clearest example of how different inducer molecules can affect antibody reactivity is the ability of T22 to detect ARA induced filaments in an ELISA, but not filaments induced by either of the polyphosphate or RNA inducers. However, T22 was reactive against RNA and polyphosphate when used in a dot blot assay. This suggests that the polyanion inducer molecules block the T22 binding, but this interaction can be disrupted through thorough wash steps that occur prior to interaction between aggregate samples and the T22 antibody. In addition, the T22 ELISA experiments also revealed that the P301S, P332S, and P364S mutations significantly decreased T22 reactive species when induced with ARA. Although the affinity of TOC and TNT antibodies does not appear to be influenced by these specific mutations, TOC does appear to show a higher affinity towards ARA induced filaments compared to fibrils induced by P100 and P700 (although the difference is not significant for P700, figure S4). Although TOC reactivity is also significantly lower for RNA induced fibrils this is most likely due to an overall lesser amount of total aggregation.

Together, these findings suggest that both the inducer molecule and disease relevant mutations can play an important role in the affinity of conformationally sensitive antibodies. In addition, different inducer molecules can provide contrasting information regarding the effects of disease relation mutations on total aggregation, filament morphology, and aggregation kinetics. Based on the data we have collected, it appears the P301S mutations has the most consistent effect on tau aggregation when comparing P to S mutations within the PGGG motif. However, other P to S mutations can also have significant effects on aggregation depending on the inducer molecule used and/or the method of detection. Until we can identify *in vitro* models that have been proven to be disease relevant, an abundance of caution should be taken when using a single inducer molecule. In particular, multiple inducer systems may be required for studies that aim to identify potential therapeutics or characterize the effects of tau aggregation conditions, disease related mutations, or post-translational modifications.

