## Supplemental Information for "*In vitro* tau aggregation inducer molecules influence the effects of *MAPT* mutations on aggregation dynamics"

*David J. Ingham[1], Kelsey M. Hillyer[1], Madison J. McGuire[1], T. Chris Gamblin[1]†*

*[1]Department of Molecular Biosciences, University of Kansas, Lawrence, Kansas, 66045, USA*

*†Department of Biology, University of Texas San Antonio, San Antonio, Texas, 78249, USA*

**Corresponding authors:**

**David J. Ingham,**

**T. Chris Gamblin,**

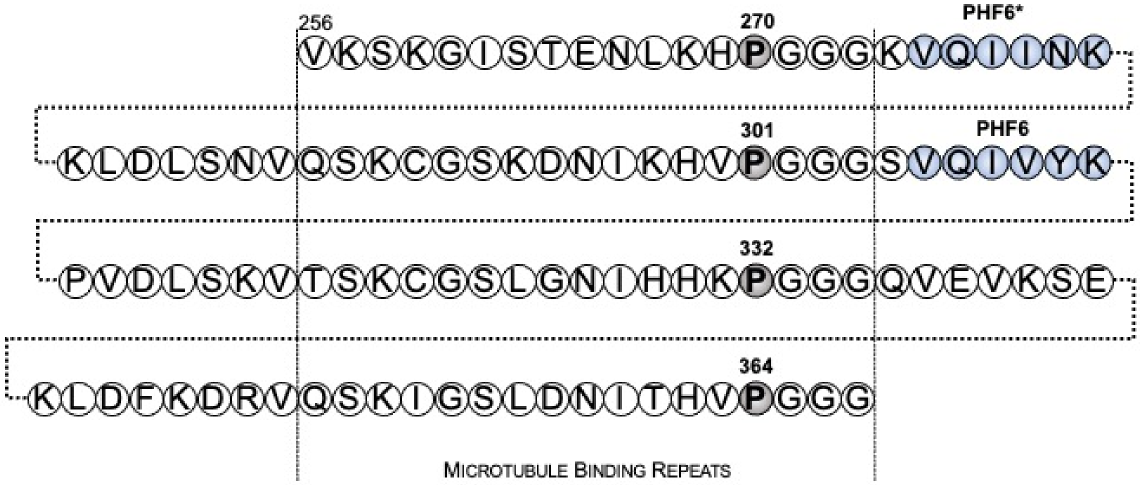


**Figure S1:** Schematic location of the PGGG motifs at the end of each microtubule binding repeat.
Motifs are found at position at the end of each microtubule binding repeat region P270 (MTBR 1), P301 (MTBR 2), P332 (MTBR 3), and P364 (MTBR 4). P270 is proceeded by the hexapeptide PHF6* motif, and P301S is proceeded by the hexapeptide PHF6, both have been demonstrated to be important in the formation of tau filaments.

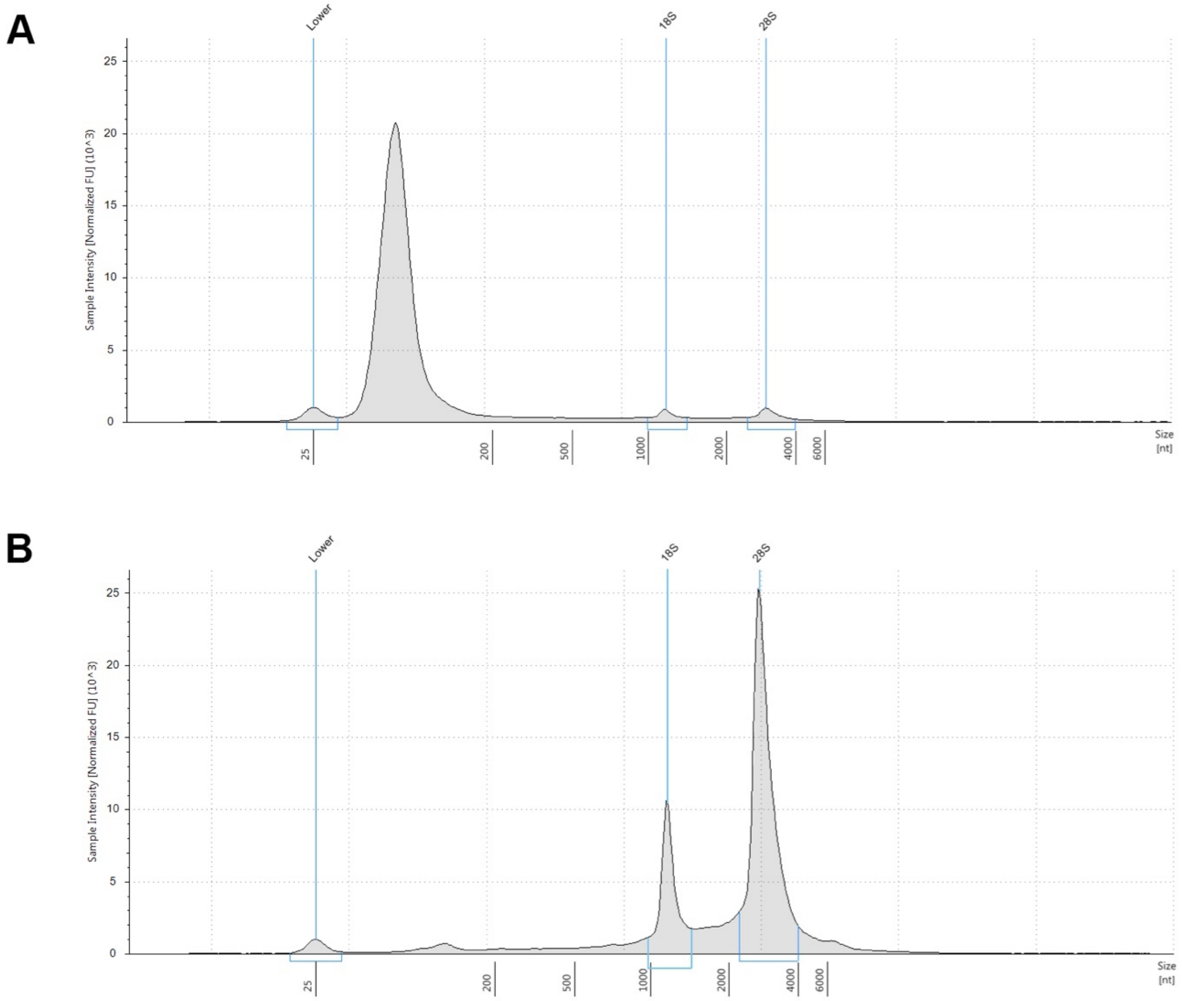


**Figure S2:** Tapestation results for A) small RNA (<200 nts), and B) long RNA (>200 nts).
Y-axis for each graph represents the band intensity based on tapestation gel image. X-axis represents the molecular weight of the band in number of nucleotides (nt). Vertical blue line labelled "Lower" at approximately 25 nt on the X-axis, represents the dye front and therefore masks any potential RNA species within the size range illustrated by the bracket on the X-axis. Vertical blue lines labelled "18S" (~900 nt -1,400 nt) and "28S" (~ 2,200 nt – 4,000 nt) indicate the expected location of 18S and 28S ribosomal RNA, respectively.

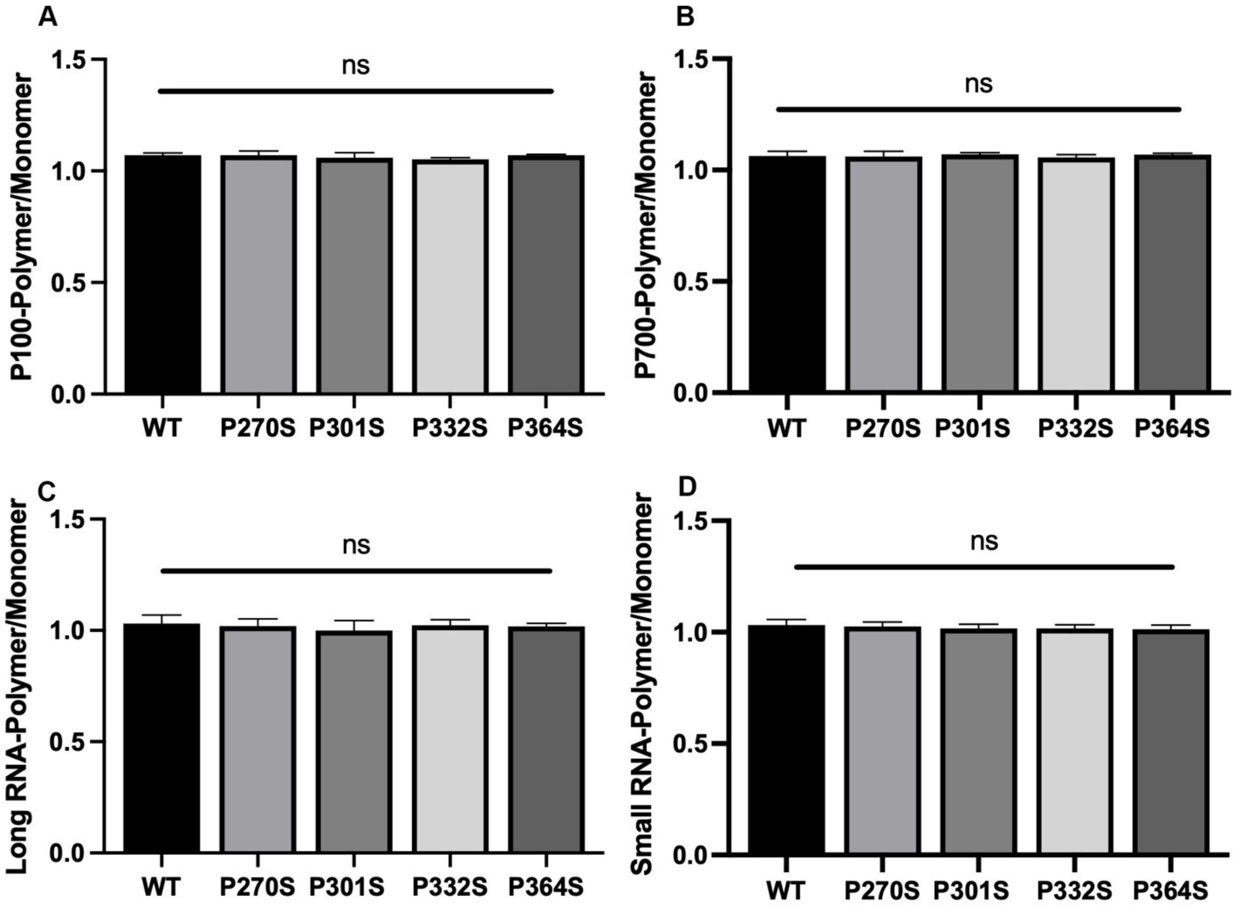


**Figure S3:** Polyphosphate and RNA 5,7,12 ELISA.
ELISA results using 5,7,12 monoclonal total-tau antibody mixture as a capture antibody. The signal for each aggregate was divided by the signal for monomeric tau (either WT or the respective P to S mutations). Y axis represents a fraction of monomeric tau signal (e.g. 1=100%). Data were analyzed using a Tukey's multiple comparison test and "ns" represents no significant difference.

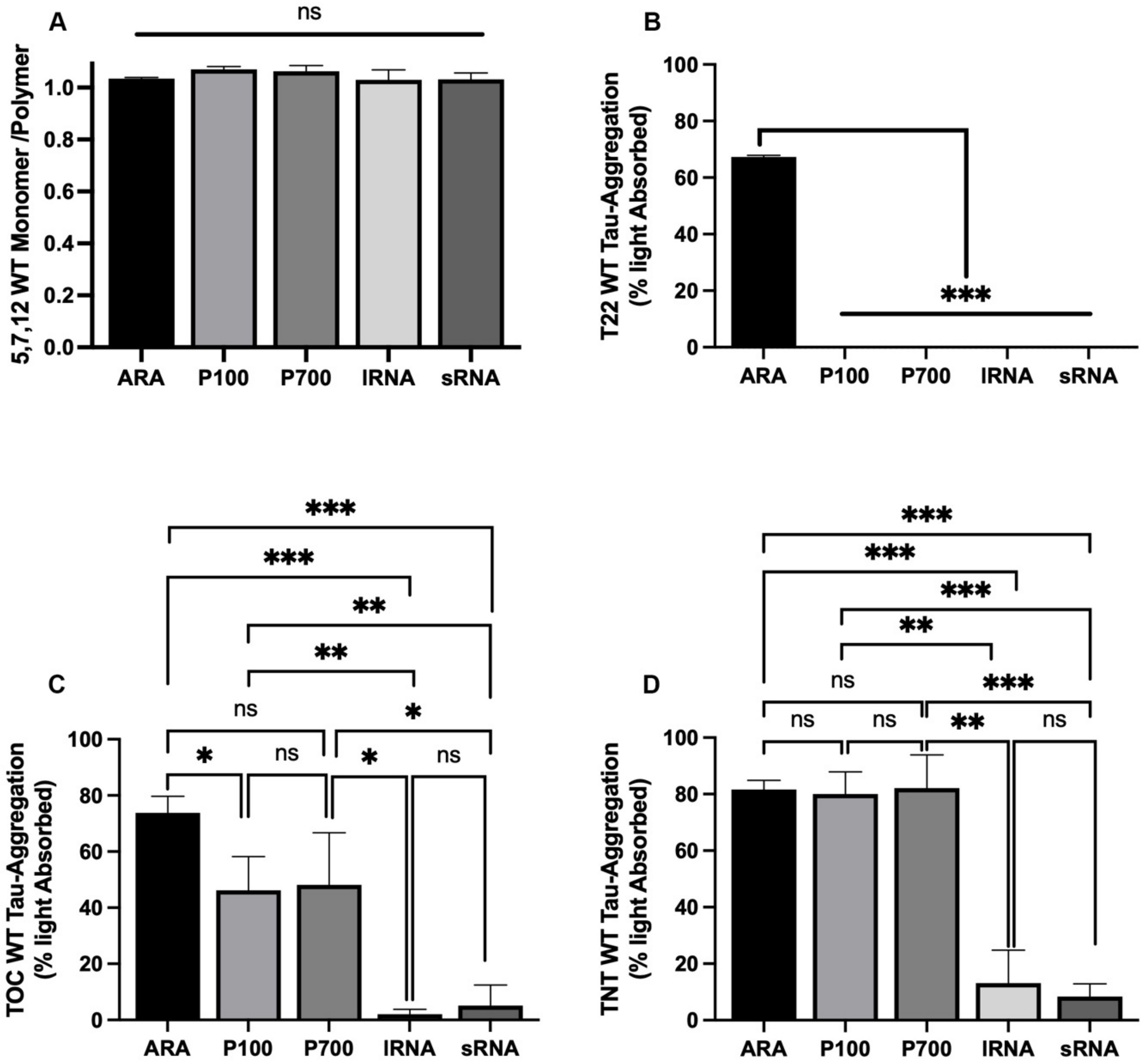


**Figure S4:** Immunoreactivity of WT HT40 using ARA, P100, P700, long RNA, and small RNA as inducer. ELISA results using 5,7,12 monoclonal total-tau antibody mixture as a capture antibody. The signal for each aggregate was divided by the signal for monomeric WT 2N4R tau. Y axis represents a fraction of monomeric tau signal (e.g. 1=100%). B-D, ELISA results of conformationally sensitive antibodies T22 (B), TOC1 (C), and TNT (D). Y-axis represents % light absorbed value (converted from A450 reading). Error bars represent SD of 3 independent experiments and data sets were compared using an un-paired t-test multiple comparison test.

$$* p \le 0.05;\ ** \ p \le 0.01;\ *** p \le 0.001.$$

*Reactivity of T22 antibody in dot-blot assay:*

Similar to filaments induced by P100 and P700 polyphosphate, neither RNA species formed T22 reactive aggregates. To ensure this was not an assay specific result, we completed a dot blot assay detecting with T22 antibody against samples of WT and P301S induced aggregates (figure 33). In this assay, polyphosphate and RNA samples did form T22 reactive species, however, although repeated experiments showed positive reactivity, the values had a high amount of variability (~50%) and therefore, we were unable to complete meaningful comparison studies.

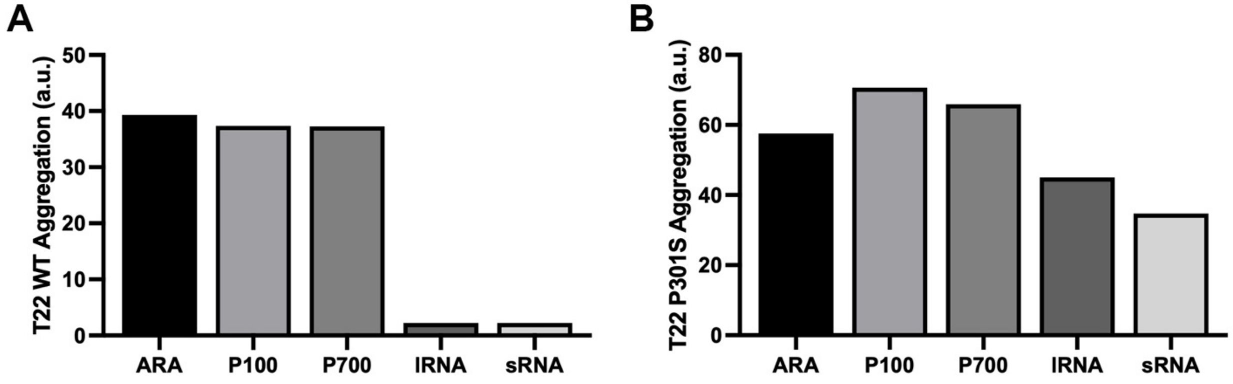


**Figure S5:** Dot-blot assay detecting endpoint aggregates using T22 antibody.
A) Comparison of WT tau induced with ARA, P100, P700, long RNA, and small RNA. B) Comparison of P301S tau induced with ARA, P100, P700, long RNA, and small RNA. Y-axis represent light intensity as measure using histogram function in Adobe Photoshop, and both data sets have been zeroed against a no inducer monomer control (n=1).

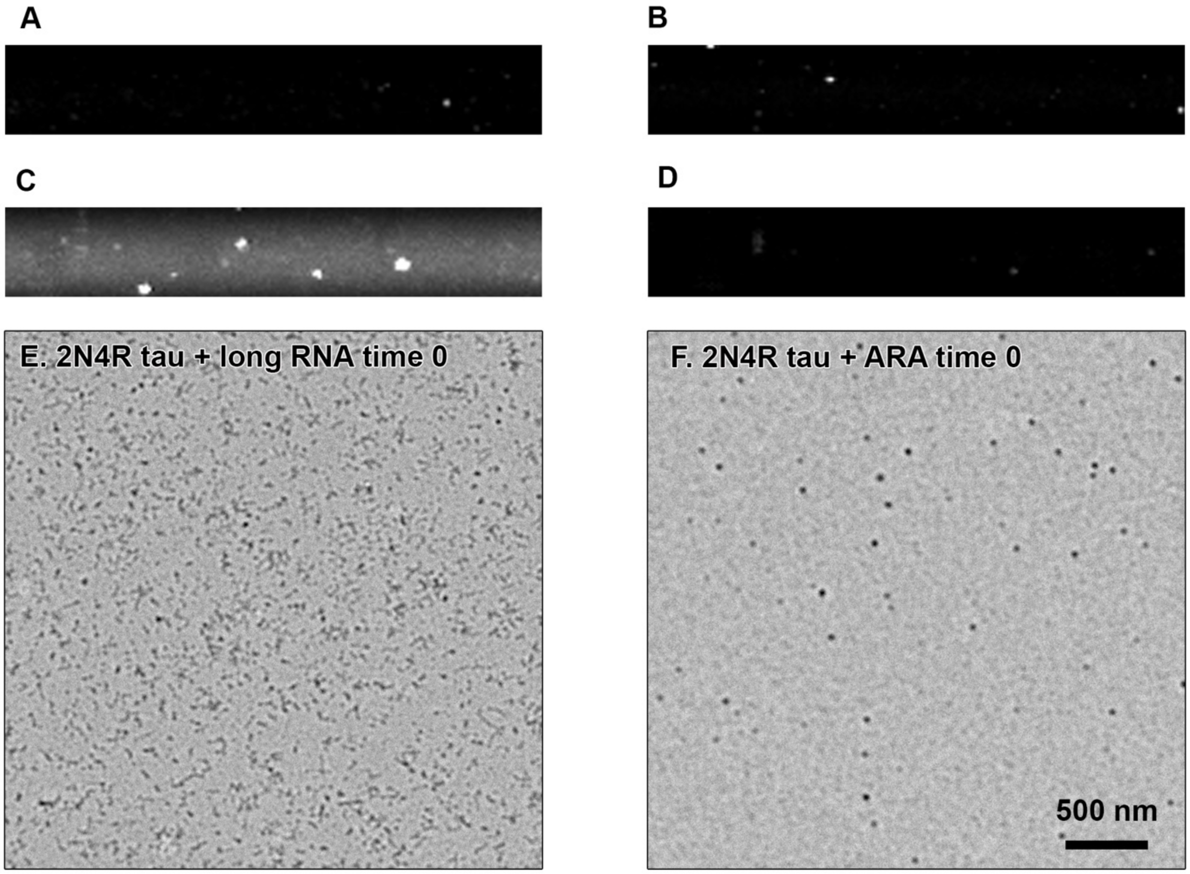


**Figure S6:** Laser light scattering and TEM upon the addition of long RNA, or ARA, to monomeric WT tau at time zero.
Right-angled laser light scattering images of rRNA only (A), 2N4R tau only (B), RNA + 2N4R tau at time zero (C) and the accompanying micrograph (E), ARA + 2N4R tau at time zero (D) and the accompanying micrograph (F). LLS images were taken at an aperture of f/5.6 and scale bar in figure F represents 500 nm for both EM micrographs.
